# Dissecting the contributions of tumor heterogeneity on metastasis at single-cell resolution

**DOI:** 10.1101/2022.08.04.502697

**Authors:** Juliane Winkler, Weilun Tan, Catherine M. M. Diadhiou, Christopher S. McGinnis, Aamna Abbasi, Saad Hasnain, Sophia Durney, Elena Atamaniuc, Daphne Superville, Leena Awni, Joyce V. Lee, Johanna H. Hinrichs, Marco Y. Hein, Michael Borja, Angela Detweiler, Su-Yang Liu, Ankitha Nanjaraj, Vaishnavi Sitarama, Hope S. Rugo, Norma Neff, Zev J. Gartner, Angela Oliveira Pisco, Andrei Goga, Spyros Darmanis, Zena Werb

**Affiliations:** Department of Anatomy, University of California, San Francisco, San Francisco CA 94143, USA; Department of Cell and Tissue Biology, University of California, San Francisco, San Francisco CA 94143, USA; Chan Zuckerberg Biohub, San Francisco CA 94143, USA; Department of Pharmaceutical Chemistry, University of California, San Francisco San Francisco CA 94143, USA; Institute of Internal Medicine D, Medical Cell Biology, University Hospital Münster, Münster, Germany; Department of Medicine, University of California, San Francisco, San Francisco CA 94143, USA; Chan Zuckerberg Biohub Investigator, San Francisco CA 94143, USA; Genentech Inc., South San Francisco, CA 94080, USA

## Abstract

Metastasis is the leading cause of cancer-related deaths, but metastasis research is challenged by limited access to patient material and a lack of experimental models that appropriately recapitulate tumor heterogeneity. Here, we analyzed single-cell transcriptomes of matched primary tumor and metastasis from patient-derived xenograft models of breast cancer, demonstrating that primary tumor and metastatic cells show profound transcriptional differences across heterogeneous tumors. While primary tumor cells upregulated several metabolic genes, metastatic cells displayed a motility phenotype in micrometastatic lesions and increased stress response signaling during metastatic progression. Additionally, we identified gene signatures that are associated with the metastatic potential and correlated with patient outcomes. Poorly metastatic primary tumors showed increased immune-regulatory control that may prevent metastasis, whereas highly metastatic primary tumors upregulated markers of epithelial-mesenchymal transition (EMT). We found that intra-tumor heterogeneity is dominated by epithelial-mesenchymal plasticity (EMP) which presented as a dynamic continuum with intermediate cell states that were characterized by novel, specific markers. These intermediate EMP markers correlated with worse patient outcomes and could serve as potential new therapeutic targets to block metastatic development.

---

Current cancer treatment is most effective in attacking the primary tumor but has little effect on metastatic cells. This is a substantial problem because metastases account for the vast majority of cancer-related deaths (1). During the multistep process of metastasis, tumor cells adapt to various microenvironments that are distinct from their site of origin, but our understanding of the processes that lead to these adaptations is limited. Moreover, phenotypic alterations of metastatic cells may also cause resistance to therapeutics that cannot be accounted for by just genotypic changes (2).

The reason why some cancers metastasize while others do not is poorly understood. For example, specific genetic alterations are not necessarily required for metastatic progression (3), highlighting the importance of phenotypic adaptations of individual tumor cells to microenvironmental influences. In order to metastasize, tumor cells have to acquire complex traits; some of these include the ability to invade, intravasate and survive in circulation until they reach the metastatic site, where tumor cells extravasate into a new tissue and give rise to a secondary tumor. One concept aiming to explain these complex phenotypic changes is that tumor cells undergo epithelial-to-mesenchymal transition (EMT) and gain mesenchymal features. Thus, EMT has been suggested to play a fundamental role for tumor cells to disseminate to distant organs (4). However, to form overt metastasis, these disseminated tumor cells need to revert the EMT process, undergo mesenchymal-to-epithelial transition (MET), and gain epithelial features again. Epithelial-mesenchymal plasticity (EMP) therefore describes the ability of tumor cells to dynamically switch between epithelial and mesenchymal cell states. EMT is often described by the loss or gain of a few canonical markers involved in cell adhesion and motility (e.g. VIM, EPCAM, CDH1, CDH2), the expression of which are regulated by a set of core transcription factors (e.g. SNAI1, SNAI2, TWIST1, ZEB1). However, these commonly used markers are context- and tissue-dependent and change dynamically during the EMT process (5–7) leading to controversies in the field that rely on these few markers (8–14). Moreover, tumor tissues are heterogeneous, displaying various phenotypes and cell states within one tumor and thus require the analysis of individual cells within one tumor. To better understand the contributions of EMP to the metastatic process we need to comprehensively analyze individual heterogeneous tumor cells both at the primary tumor and metastatic site.

Advances in single-cell transcriptomics have enabled investigation into intra-tumor heterogeneity in breast cancer (BC) (15–18) and many other cancers (19–21). For instance, an integration of these studies across multiple different tumor entities has highlighted both the importance and the context-dependency of the EMT process in tumor biology (22). However, with a few notable exceptions (23, 24), these studies focus on characterizing primary tumors. Moreover, they lack information about patient outcomes and metastatic phenotypes due to the necessary long-term follow-up. Comparing tumor heterogeneity at single-cell resolution between matched primary and metastatic tumors is logistically difficult; patient metastatic tumor samples are often collected years after the primary tumor was resected. Moreover, analyzing metastatic lesions is also technically difficult because they may consist of individual or small numbers of metastatic cells within complex tissues, which are hard to locate and isolate from patients. It is particularly challenging to investigate EMP *in vivo*, in both primary tumors and metastatic lesions, using an unperturbed system that resamples heterogeneous human tumor tissue. Finally, while xenograft models of metastasis can alleviate many of these limitations, such models that rely on transplanted cell lines do not faithfully reproduce the heterogeneity present in primary tumors. Thus, a detailed understanding of the involvement of the dynamic and context-dependent EMP process in metastasis is lacking (14).

Here, we characterized the metastatic potential of a large panel of patient-derived xenograft (PDX) models of human BC that spontaneously metastasize and preserve the heterogeneity of the primary human tumor. We analyzed the transcriptional profiles of individual primary tumor and matched metastatic cells using different single-cell RNA sequencing (scRNA-seq) approaches. Our study provides a rich dataset that allows us to investigate the impact of tumor heterogeneity on metastatic phenotypes both at the primary tumor and metastatic site at single-cell resolution. We identified gene signatures that are associated with metastatic potential. Specifically, we found that highly metastatic tumors express elevated EMT markers and demonstrate that EMP is a key factor of intra-tumor heterogeneity both at the primary tumor and the metastatic site. Within the continuum of EMP, we identified intermediate EMP cell states that are characterized by specific marker genes. High expression of those EMP marker genes was correlated with worse outcomes in a subset of BC patients.

## BC PDX models with varying metastatic potential show transcriptional heterogeneity

To investigate intrinsic factors that impact a tumor’s ability to metastasize, we analyzed the transcriptional heterogeneity of primary tumors as well as matched metastases at single-cell resolution using PDX BC models (Figure 1A). Human breast tumors were orthotopically transplanted into the cleared mouse mammary fat pad and spontaneously metastasized to the lung and other organs. They thereby preserve the heterogeneity of the primary human tumor, fully recapitulate the metastatic cascade, and mimic the metastatic pattern of the patient (25–27). We characterized PDX models derived from 13 BC patients, belonging to different BC subtypes (three luminal B, ten basal) with varying metastatic potential. Our PDXs included two estrogen receptor (ER) and progesterone receptor (PR) positive, one triple-positive (ER, PR, HER2), and ten triple-negative BC (TNBC) models (25, 28); three of the basal TNBC PDX models were newly established in this study (Supplementary Table 1).

**Figure 1.**
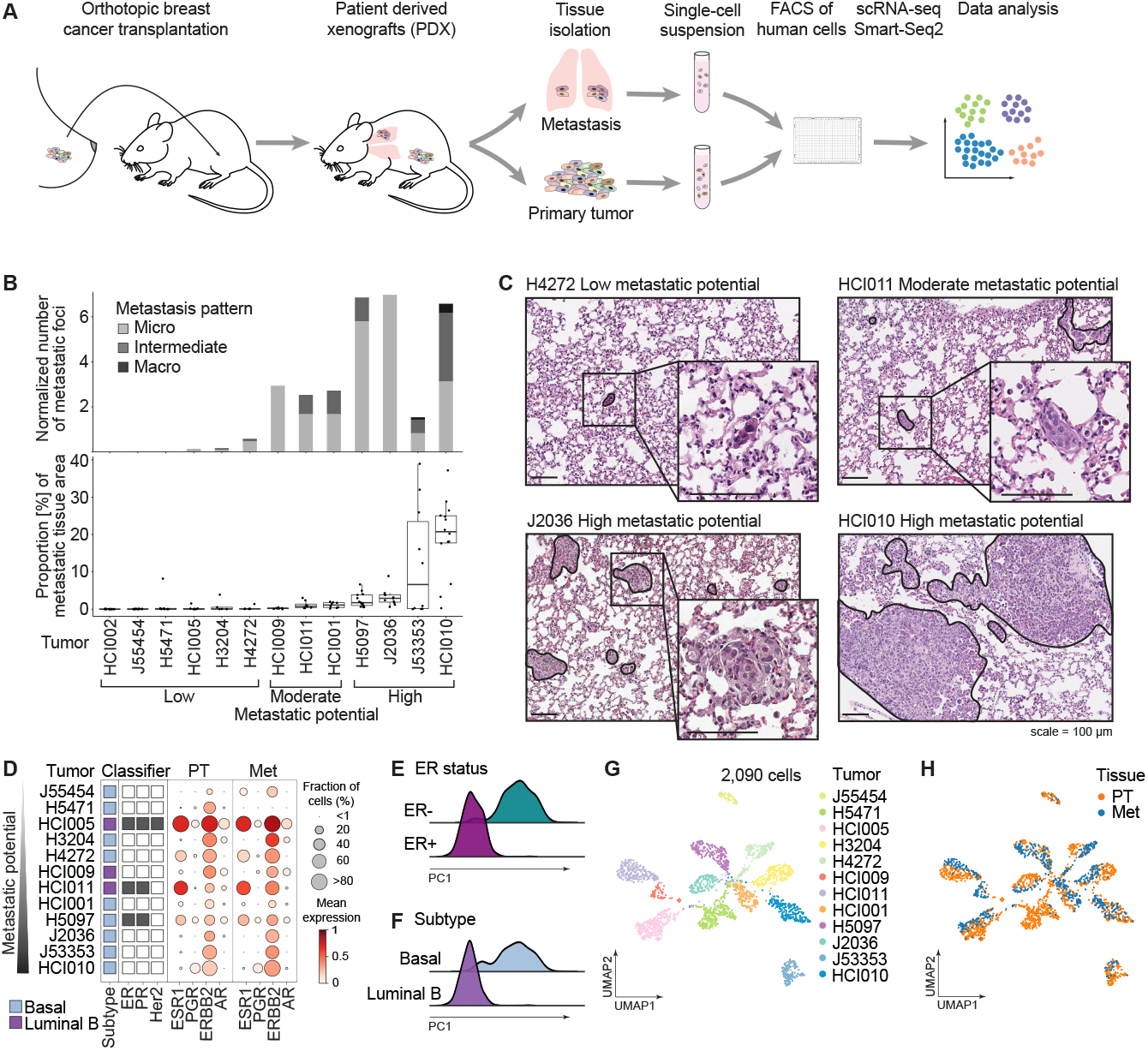
BC PDX models with varying metastatic potential show transcriptional heterogeneity. (**A**) Schematic overview of the experimental setup. Metastatic lung and primary tumor tissue were isolated from BC PDX models and dissociated. The resulting single-cell suspensions were enriched for human cells, sorted into 384 well plates and scRNA-Seq was performed using Smart-Seq2. (**B**) Bar chart shows the median number of metastatic foci per mm^2^ lung tissue area per tumor model (upper panel) determined by histology. The size of metastatic foci is colored in shades of gray (micrometastasis: < 10 cells, intermediate: 10–100 cells and macrometastasis: > 100 cells). Boxplot shows the fraction of metastatic tissue per total lung tissue area determined by histology. Annotations indicate the metastatic potential of the tumor models. (**C**) Representative H&E images of metastatic lung tissues of tumor models for low, moderate and high metastatic potential. Scale = 100 µm. (**D**) Bubble plot shows the expression of receptors in primary tumor (PT) and metastatic cells (Met) per tumor model. The size of dots indicates the fraction of expressing cells and the red color indicates the magnitude of gene expression. Box annotations show BC subtype and receptor classification. Tumor models are ordered by increasing metastatic potential as determined in (B). (**E**) Ridgeplot shows the normalized number of cells along Principal Component 1 (PC) coordinates color-coded by ER status. (**F**) Ridgeplot shows the normalized number of cells along PC1 coordinates color-coded by BC subtype. (**G**) UMAP projection of single-cell transcriptomes color-coded by individual tumor models. (**H**) UMAP projection of single-cell transcriptomes color-coded by primary tumor (PT, orange) and metastatic cells (Met, blue).

First, we characterized the metastatic phenotype of the different tumor models once primary tumors reached a size of 2.5 cm in diameter. Based on the number and size of metastatic foci in the lungs of recipient mice the tumor models were grouped into those with low (n=6 models), moderate (n=3), and high metastatic potential (n=4). PDX models with low metastatic potential form no or very few micrometastases (< 10 cells), moderate models show more micro- and intermediate-sized (10 - 100 cells) metastases, and highly metastatic models develop either a high number of micrometastases and/or many macrometastases (> 100 cells) resulting in a substantial metastatic burden (proportion of metastatic cells in the lung) (Figure 1B, C, Supplementary Figure S1A, B). The metastatic potential based on this classification was independent of the primary tumor growth rate (Supplementary Figure S1C). For example, the fast-growing but poorly metastatic HCI002 model developed very few but larger metastatic foci even after primary tumor resection with subsequent tumor recurrence (Supplementary Figure S1E, F). Tumor resection allowed HCI002 to grow for a similar period as the slower growing but highly metastatic HCI010 model (Supplementary Figure S1D), indicating that HCI002’s low metastatic potential is independent of primary tumor growth rate.

To investigate the transcriptional landscape of primary tumor and metastatic cells, individual tumor cells were isolated from primary tumors and matched metastatic lungs from 12 PDX models for scRNA-seq. Tumor cells stained with a human-specific antibody directed against a ubiquitous cell surface marker (CD298) (29) were isolated by fluorescent activated cell sorting (FACS) and subjected to scRNA-seq (Smart-Seq2). High-quality single-cell transcriptome data were collected for 2,090 cells (1,395 primary tumor and 695 metastatic cells). Of note, we were not able to isolate a sufficient number of metastatic cells from the poorly metastatic HCI002 model. The PAM50 BC subtype (Supplementary Table 1) and receptor status was confirmed for most samples (Figure 1D) according to ESR1 (ER), PGR (PR), and ERBB2 (HER2) transcript detection. Interestingly, ERBB2 was detected in all tumors including those not clinically classified as HER2-positive. This was potentially due to the required threshold for the clinical classification of the original tumor by histochemistry and/or single region sampling of the heterogeneous original tumor. In addition, receptor expression was maintained in metastatic cells in our data (Figure 1D and Supplementary Figure S1G). These results are in contrast to studies that reported a change of receptor status during tumor progression and recurrence occurring in up to 40% of patients depending on the specific receptor, which had implications for treatment options and poor patient outcomes (30–32). However, our data indicate that changes in receptor status are not caused by the metastatic process but are likely a consequence of selection during receptor-targeted therapy.

The ER status and BC subtype were the major sources of variation in our dataset. This is illustrated by principal component (PC) analysis, which showed a clear separation of ER status and BC subtypes along PC1 (Figure 1E, F, Supplementary Figure S1H). Moreover, individual tumors clustered separately from other tumors, reflecting the effect of interpatient heterogeneity on gene expression (Figure 1G). Notably, variability between technical batches (individual plates) or biological replicates (same tumor implanted into different animals) was not observed (Supplementary Figure S1I, J). Finally, within individual tumor models, primary and metastatic cells clustered separately in all cases, highlighting the transcriptional differences between primary and metastatic cells from a particular tumor model (Figure 1H).

Taken together, we established and characterized PDX models of different BC subtypes with varying metastatic potentials that were independent of their primary tumor growth rate. Receptor status was maintained between primary tumor and metastatic cells. In addition to interpatient heterogeneity, primary tumor and metastatic cells showed strong transcriptional differences within individual tumors.

## Differential gene expression analysis reveals metastasis-associated gene signatures and heterogeneity between cells

To characterize general transcriptional programs unique to metastatic cells, cells were grouped across all samples by tissue source (primary tumor or metastatic lung) and differential gene expression was determined using MAST (33) with the tumor model as a covariate. We found 132 differentially expressed genes (DEGs), 79 of which were upregulated in metastatic cells conserved across all 12 tumor models (log_2_ fold change > 0.5; Figure 2A, Supplementary Figure S2A, Supplementary Table 2). Among the top metastasis-associated genes were several cytokeratins (KRT5, KRT6B, KRT14, KRT17, KRT81), calcium-binding S100 proteins (S100A16, S100A14), heat shock protein HSP1, cell-surface proteins such as TSPAN1, serine proteases (KLK6, KLK7), and the glycoproteins CEACAM6 and PSCA. Pathway-level analysis revealed that metastatic cells were enriched in MYC, E2F, PI3K/AKT/MTOR signaling and oxidative phosphorylation (Figure 2B). This observation is consistent with studies showing enrichment of MYC signaling and oxidative phosphorylation pathways in metastatic BC cells found in the lung (29, 34). Interestingly, metastatic cells additionally upregulated genes involved in immune response pathways (IL6/JAK/STAT3), presumably as an adaptation to the metastatic microenvironment (Figure 2B, Supplementary Figure S2B). In contrast, hypoxia, EMT, angiogenesis, and glycolysis were pathways enriched in primary tumor cells (Figure 2B). To determine whether the identified pathways were upregulated in all individual primary tumor or metastatic cells or only in a subset of cells we examined the expression of DEGs associated with the top enriched pathways in either primary tumor (hypoxia) or metastatic cells (MYC). The analysis revealed a profound heterogeneity both between and within tumor models (Supplementary Figure S2C). However, when analyzed individually, primary tumor and metastatic cells displayed strong transcriptional differences illustrated by separation along PC2 (Figure 2C; Supplementary Figure S2D–G) mirroring our previous observations (Figure 1H).

**Figure 2.**
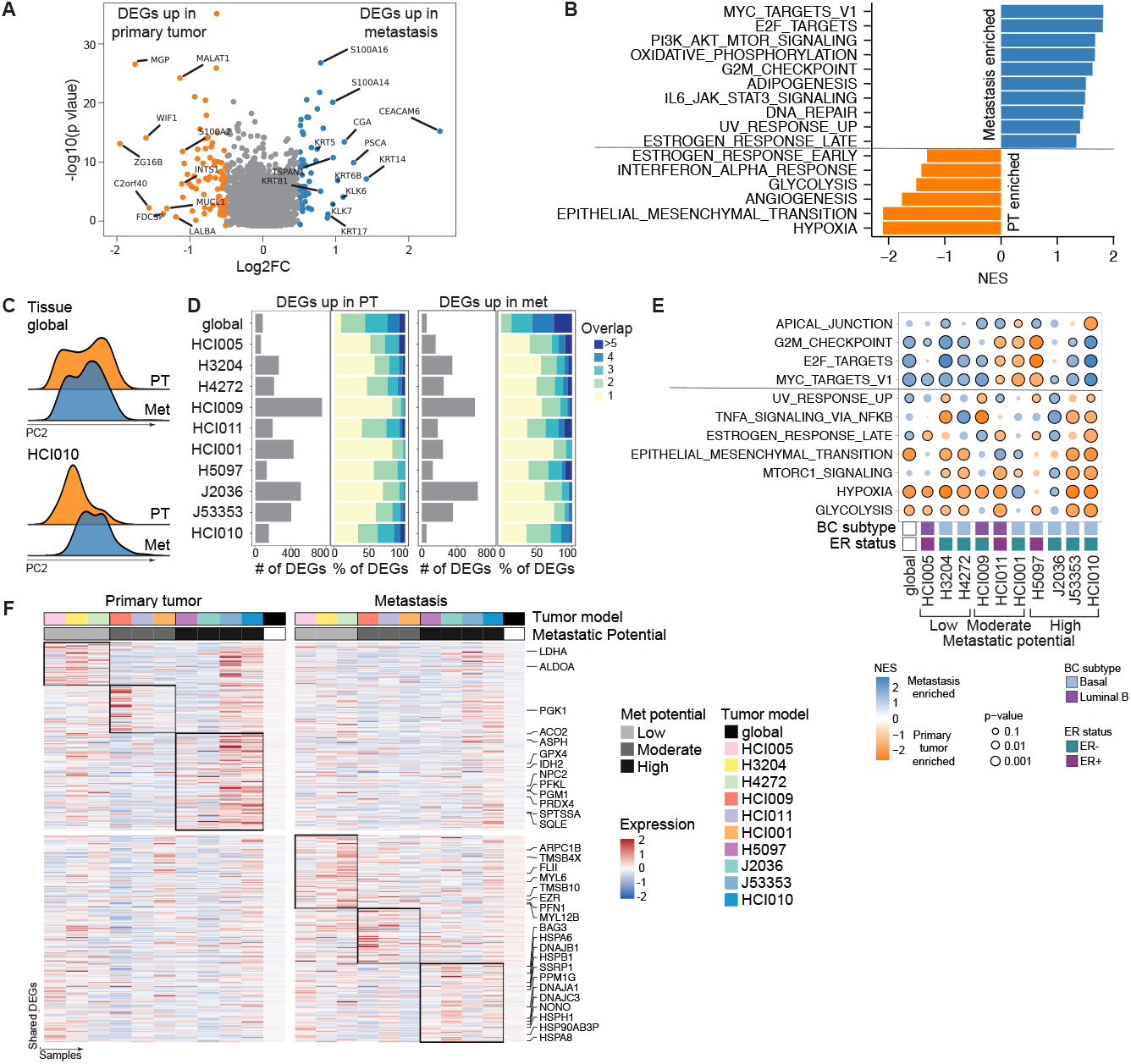
Differential gene expression between primary tumor and matched metastatic cells. (**A**) Volcano plot shows log2 fold change of expression and p-value of DEGs in primary tumors vs metastases. Highlighted are top DEGs. (**B**) Barplot shows pathways enriched in primary tumors (negative normalized enrichment score (NES), orange) and metastases (positive NES, blue) using HALLMARK gene sets from MSigDB. (**C**) Ridgeplots show normalized cell counts along PC2 color-coded by primary tumor and metastatic cells of all tumor models grouped together (global, upper panel) and a representative individual tumor model (HCI010, lower panel). (**D**) Bar charts show the number of DEGs (gray bars) upregulated in primary tumors (PT, left) and metastases (met, right) for each individual tumor model. Color bars indicate the proportion of DEGs that are shared between two or multiple individual tumor models (blue color scale) or exclusive to one tumor model (yellow). (**E**) Bubble plot shows enriched HALLMARK pathways (MSigDB) that are shared between at least four tumors. NES indicates enrichment in primary tumors (negative, orange) or metastasss (positive, blue), size of bubbles indicates p-values, and circled outline indicates significant p-value < 0.05. (**F**) Heatmaps show mean expression in individual tumor models of DEGs upregulated in the primary tumor (left) or metastasis (right) that are shared between at least 2 tumor models within the same metastatic potential group (black box). Annotations indicate selected DEGs, tumor model, and metastatic potential.

To control for this pronounced variability amongst our tumor models, we next analyzed DEGs between primary tumor and metastatic cells for each model separately (Supplementary Table 2) and compared these across tumor models. Due to insufficient metastatic cell numbers, two tumor models with low metastatic potential (J55454, H5471) were excluded from this analysis (Supplementary Figure S2E, F). The different tumor models showed a wide range of numbers of DEGs (Figure 2D). Notably, more than 50% of DEGs were tumor model-specific and only a few (< 5%) were shared between more than 5 tumor models, highlighting again the magnitude of inter-patient heterogeneity (Figure 2D). We focused on enriched pathways that were shared between tumor models (Figure 2E). Although most shared pathways were also identified in the previous analysis across tumor models (Figure 2B, Supplementary Figure S2B), some pathways showed intriguing enrichment differences between tumor models. For example, whereas the combined analysis revealed an overall suppression of the estrogen-response pathway in the primary tumor, the individual analyses showed that this pathway was specifically upregulated in ER+ primary tumors (HCI005, HCI011, H5097) compared to matched metastatic samples. This suggests that estrogen signaling is impaired in the metastatic cells despite maintained ESR1 expression (Figure 2E, Figure 1D, Supplementary Figure S1G). Additionally, while this analysis showed that metastatic cells of some tumor models were enriched in the G2M checkpoint pathway, we could not confirm an overall more active proliferation or substantial cell cycle shifts of metastatic cells in our data (Supplementary Figure S2H, I). Owing to their larger size, primary tumors have limited access to nutrients; thus, it is not surprising that enrichment of glycolysis and hypoxia seemed to be a general feature in primary tumors. Moreover, EMT was enriched either in primary tumors or metastasis in the majority of the analyzed tumor models indicating a dynamic activity of this pathway in both compartments.

Since individual DEGs were shared only between a few tumor models, we focused on DEGs that were common between tumor models with a similar metastatic phenotype (Figure 2F, Supplementary Table 3). We found 74 upregulated genes in metastatic cells that were shared between at least two tumors of low metastatic potential. Among these were many genes involved in cytoskeleton assembly and cell motility (e.g. MYL12B, MYL6, PFN1, TMSB4X, TMSB10, ARPC1B, EZR, FLII). In contrast, among the 91 genes upregulated in metastatic cells from high metastatic tumor models were many genes indicative of high stress-response signaling, including several heat shock proteins (HSPB1, HSPA8, HSPA6, HSPH1, HSP90AB3P, DnaJs A1, B1, C3, and BAG3), PPM1G and genes involved in DNA damage repair (SSRP1, NONO). Several genes involved in glycolysis (ALDOA, LDHA, PGK1, PFKL, PGM1) and other metabolic processes (GPX4, PRDX4, ACO2, ASPH, IDH2, SQLE, NPC2, SPTSSA) were upregulated in primary tumor cells suggesting differential metabolism in primary tumors as compared to the metastasis.

In summary, we observed strong transcriptional differences between primary tumor and metastatic cells on an individual tumor level with the majority of DEGs being specific to each tumor. Shared features across models are upregulation of hypoxia, glycolysis and other metabolic-related genes in primary tumor cells. Shared upregulated genes among metastatic cells are involved in cytoskeleton assembly and motility and stress response signaling.

## Metastatic signatures are correlated with patient outcomes

The tumor models used in this study exhibit consistent metastatic behaviors and were classified into tumors with low, moderate, and high metastatic potential (Figure 1B, C, Supplementary Figure S1A). One fundamental question is whether intrinsic features of the primary tumor are predictive of the observed different metastatic potential of those tumor models. To address this question we generated an additional, larger scRNA-Seq dataset, which better reflected the intra-tumor heterogeneity of the primary tumors. To this end, we performed high-throughput, droplet-based scRNA-Seq with MULTI-Seq (36) sample multiplexing on 10 different primary tumors with varying metastatic potential (Figure 3A), resulting in 16,861 tumor cells (Supplementary Figure S3A–C).

**Figure 3.**
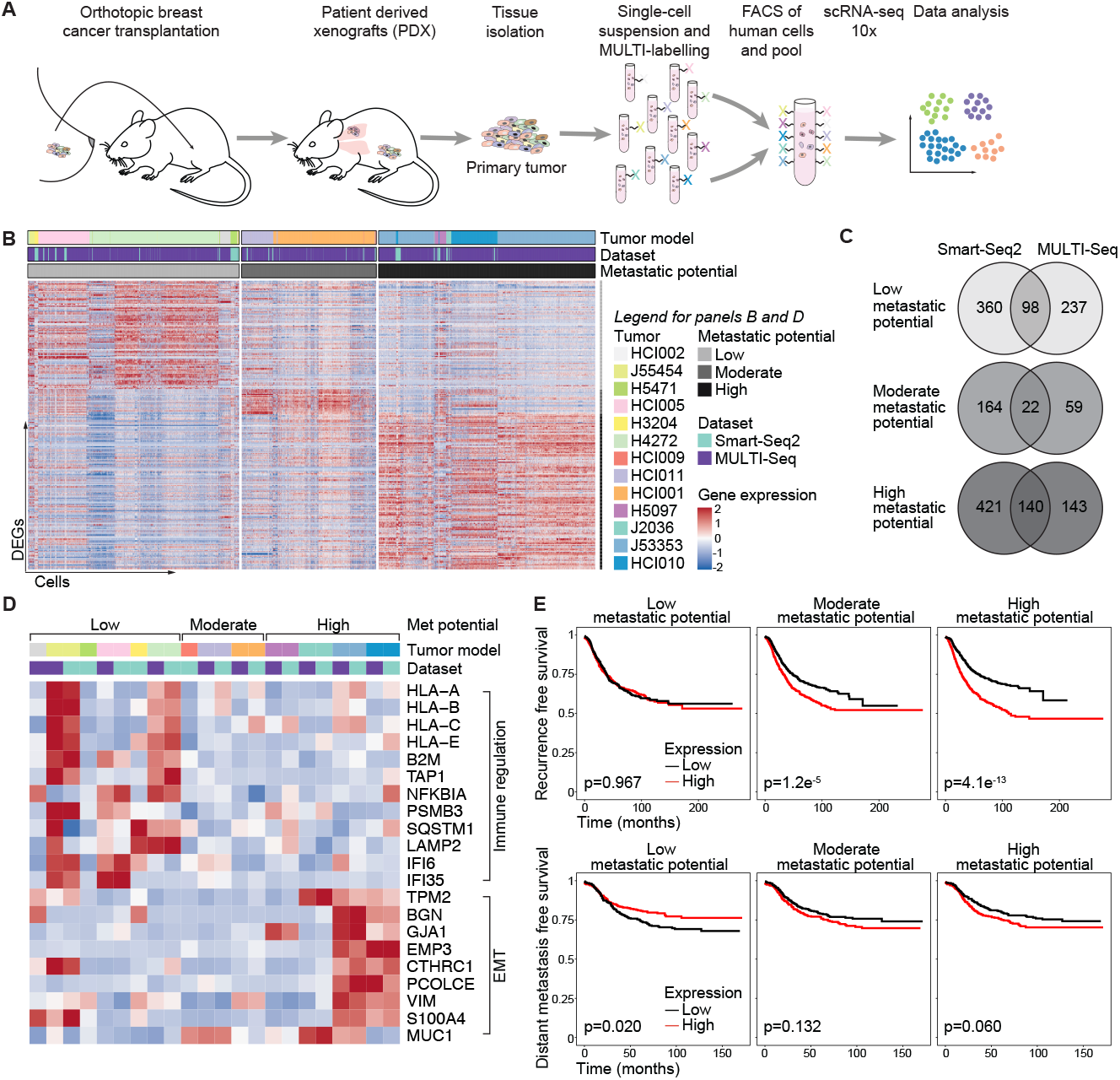
Metastatic signatures are correlated with patient outcomes. (**A**) Schematic workflow about the experimental setup using MULTI-Seq. (**B**) Heatmaps show DEGs between individual tumors and tumors of the other metastatic potential groups that are shared between at least 2 tumors. Annotations show the tumor model, dataset and metastatic potential group. Venn diagram shows the number of DEGs shared between the Smart-Seq2 and MULTI-Seq datasets for the different metastatic potential groups. (**D**) Heatmaps show the mean expression of selected metastasis-associated genes. Legends for annotations are the same as in Figure 4B. (**E**) Recurrence-free survival (RFS, top, n = 2,032 patients) and distant metastasis-free survival (DMFS, bottom, n = 958 patients) of BC patients using the mean expression of the metastasis-associated gene signatures (generated with KM-plotter (35)).

To identify signatures that were associated with the metastatic potential of the primary tumor, we looked for DEGs between different metastatic potential groups (Figure 3B, Supplementary Table 4, Supplementary Table 5). For each metastatic potential group, we selected genes that were shared between both scRNA-Seq methods that were used in this study (Figure 3C, Supplementary Table 6). Among the shared genes upregulated in primary tumors with a low metastatic potential were genes related to immune regulation processes such as antigen processing and cross-presentation (e.g. HLA-A, HLA-B, HLA-C, HLA-E, B2M, TAP1), and innate immunity (e.g. NFKBIA, PSMB3, SQSTM1, LAMP2, IFI6, IFI35) (Figure 3D, Supplementary Figure S3D). As our model used immunocompromised mice that lack B, T and NK cells, these findings potentially reflect a tumor-intrinsic, anti-metastatic feature independent of the canonical function of these genes in immune regulation. Genes upregulated in highly metastatic primary tumors included known metastasis-related genes such as S100A4 (37–39), MUC1 (40) and genes associated with EMT (VIM, PLOD1, BGN), including the common EMT marker vimentin (VIM) (Figure 3D). MYC signaling was among the top 5 enriched pathways in highly metastatic primary tumors (Supplementary Figure S3E). MYC signaling can lead to evasion from immune surveillance by the suppression of interferon signaling and antigen-presentation pathways including the down-regulation of B2M and MHC-I (41, 42). This anti-correlation could explain the observed upregulation of immune regulatory pathways in poorly metastatic compared to highly metastatic primary tumors that showed elevated MYC signaling (Supplementary Figure S3F). Supporting our experimental data (Supplementary Figure S1C) a highly metastatic phenotype is not the result of more proliferation since proliferation rate or cell cycle phase distributions were not significantly changed between primary tumors of different metastatic potentials (Supplementary Figure S3G, H).

Next, we tested whether the observed metastasis-associated signatures were correlated with patient-related outcomes using publicly available bulk gene expression data of BC patients across different subtypes (35) (Figure 3E). Indeed, patients with a high expression of the poorly metastatic signature exhibit improved distant metastasis-free survival (DMFS). A high expression of moderate metastatic genes was associated with worse recurrence-free survival (RFS) and a high expression of the highly metastatic signature showed the worst outcome for patients.

In summary, we identified intrinsic metastasis-associated gene signatures in primary tumors that were correlated with patient-related outcomes of an external dataset. While genes upregulated in poorly metastatic primary tumors are involved in immune regulation presenting potential non-canonical anti-metastatic functions, genes present in the highly metastatic signature were associated with EMT.

## Epithelial-mesenchymal plasticity is a key feature of tumor heterogeneity and is associated with metastatic potential

Markers of EMT were upregulated in primary tumors of highly metastatic tumor models as compared to models with low metastatic potential. However, we also found EMT to be enriched in either primary tumor or metastatic cells in different tumor models indicating a dynamic process during metastatic progression. Tumor cells must switch phenotypes multiple times during the metastatic process to adapt to different environments. While numerous studies have established a role for the epithelial-mesenchymal transition (EMT) in cancer progression, fewer have examined the role of epithelial-mesenchymal plasticity (EMP) in this process (14). Studying the latter is challenging, as most studies focus on a limited set of end-point markers to distinguish epithelial and mesenchymal cell states and/or to perturb these markers to test the role of EMT either *in vitro* or in mouse models *in vivo*. Here, using single-cell transcriptomics, we seek to identify those cell states across the spectrum of EMT and in multiple heterogeneous human tumor populations that correlate with their metastatic potential *in vivo*. To this end, we used a pan-cancer gene signature of 303 mesenchymal and epithelial markers to characterize the EMP state of individual cells (43). Individual canonical epithelial markers (EPCAM and CDH1) were highly expressed in cells with a high epithelial signature and also mesenchymal markers (VIM, FN1, CDH2) showed the expected expression patterns (Supplementary Figure S4A). However, some of these commonly used markers, such as FN1 and CDH2, were minimally detected on an individual cell level, indicating the importance of using multi-gene signatures to define cell states. To illustrate that cells can express epithelial and mesenchymal markers dynamically, we combined epithelial and mesenchymal signatures to define the overall EMP cell state; e.g. an EMP signature >0 reflects cells with a higher mesenchymal signature than epithelial signature. These EMP signatures of individual tumor models were strongly correlated (R^2^ = 0.78) between our two datasets (Smart-Seq2 and MULTI-Seq), demonstrating reproducibility of results across different sequencing methods and experiments (Supplementary Figure S4B). Tumors that consistently metastasized expressed a significantly higher (p<0.001) EMP signature compared to those that poorly metastasized (Figure 4A, Supplementary Figure S4C). For individual tumor models, a high EMP state was associated with metastatic potential (Figure 4B, Smart-Seq2 R = 0.336, Supplementary Figure S4D, MULTI-Seq R = 0.606). Surprisingly, the overall EMP state of each tumor model was similar for both primary tumor and metastatic cells (Figure 4C) suggesting an intrinsic determinant of EMP that is potentially independent of environmental influences which were likely very different between the tissues. However, across individual cells, the EMP state was highly variable within one tumor model. Indeed, EMP signatures of individual tumor models were strongly correlated with PC1 coordinates, indicating that the EMP cell state is a major source of variation between cells within one tumor model and significantly contributes to intra-tumor heterogeneity (Supplementary Figure S4 F). Finally, we observed that EMP state was gradually changing in transcriptional space, further illustrating that EMP is a continuum of cell states (Figure 4 D).

**Figure 4.**
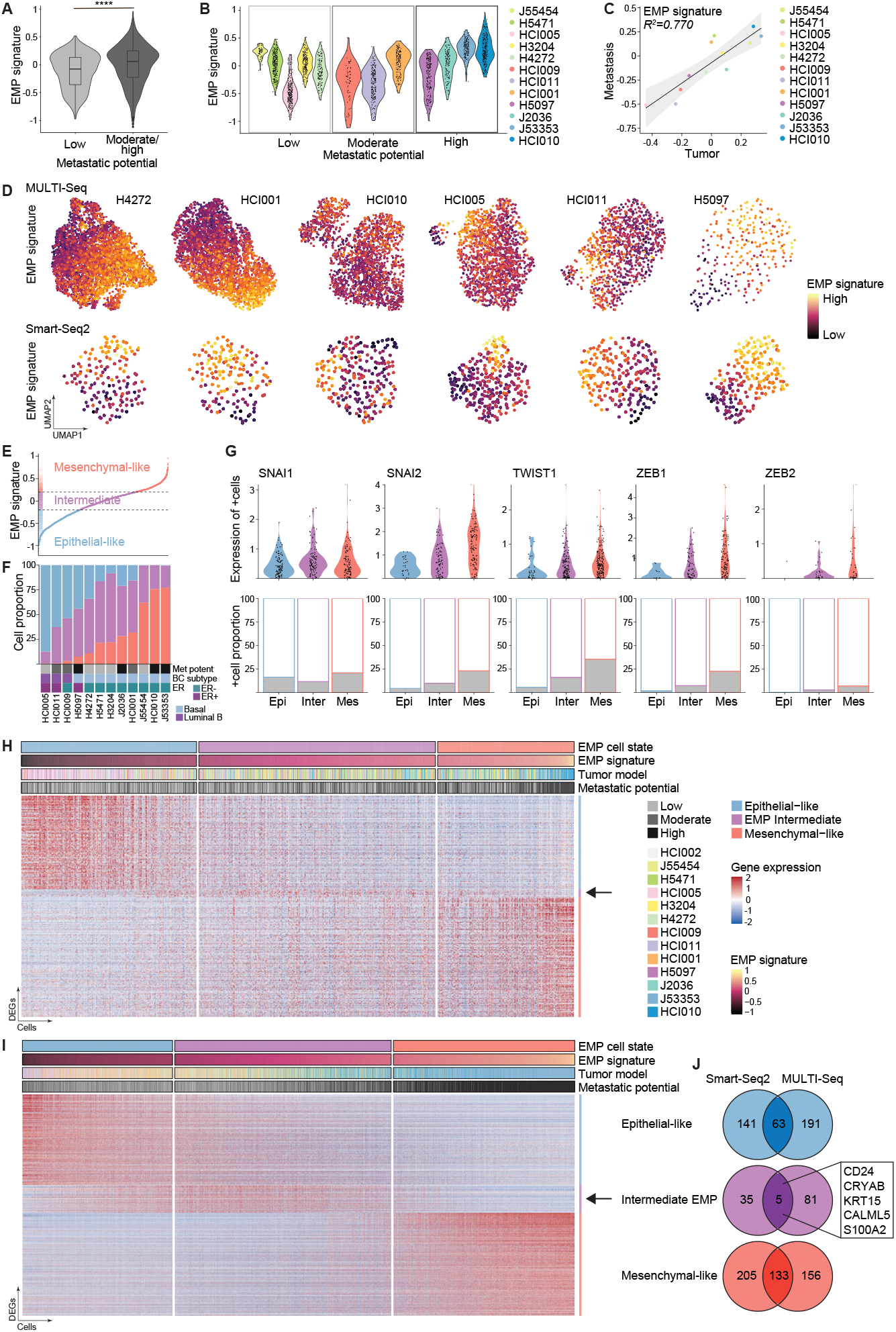
EMP is a key feature of tumor heterogeneity. (**A**) Violin plot shows EMP signature expression of tumor models with low and intermediate/high metastatic potential using the Smart-Seq2 dataset. Boxplot showing median, significance p<0.001 by Wilcox test. (**B**) Violin plot shows EMP signature per tumor model ordered by metastatic potential using the Smart-Seq2 dataset. (**C**) Scatter plot shows the correlation of the mean EMP signature of the primary tumor and metastatic cells colored by the tumor model. Linear regression with 95% confidence intervals and Pearson correlation coefficient are shown. UMAP projections of single-cell transcriptomes for individual tumor models. The color scale indicates the magnitude of EMP signature expression. (**E**) Cells ranked by EMP signature define three cell states: epithelial-like (blue), intermediate EMP (purple) and mesenchymal-like cells (red) using the Smart-Seq2 dataset. (**F**) Bar chart shows the proportion of the three different EMP cell states in each tumor model ranked by the increasing proportion of mesenchymal-like cells. Gray-scale boxes indicate the metastatic potential. Other annotations indicate ER status and BC subtype. Showing the Smart-Seq2 dataset. (**G**) Violin plots (top) show expression of EMT-associated TFs in expressing cells grouped by EMP cell states (Epi = epithelial-like, Inter = Intermediate EMP, Mes = mesenchymal-like cells). Bar charts (bottom) show the fraction of expressing cells colored in gray. Showing the Smart-Seq2 dataset. (**H**) Heatmap shows DEGs for epithelial-like, mesenchymal-like, and intermediate EMP cells for the Smart-Seq2 data. Cells are ordered by increasing EMP signature. Annotations indicate EMP cell state, EMP signature expression, tumor model and metastatic potential. The arrow highlights intermediate EMP cell marker genes. (**I**) Same as in (H) using the MULTI-Seq data. (**J**) Venn diagrams show overlap DEGs of epithelial-like, mesenchymal-like, and intermediate EMP cells between Smart-Seq2 and MULTI-Seq data. Highlighted are overlapped markers for intermediate EMP cells.

Next, we asked whether the metastatic potential is associated with the EMP state of individual cells or their proportion within the tumor. Cells were classified by the magnitude of their EMP signature expression into three different cell states: epithelial-like (epithelial > mesenchymal signature), intermediate EMP (epithelial = mesenchymal signature) and mesenchymal-like cells (epithelial < mesenchymal signature) (Figure 4E Supplementary Figure S4G The proportion of mesenchymal-like cells largely aligned with the metastatic potential, with no classified mesenchymal-like cells present in poorly metastatic tumors and almost no epithelial-like cells present in tumors with high metastatic potential (Figure 4F, Supplementary Figure S4H ER+ and luminal B tumors showed the highest proportion of epithelial-like cells. However, even within this group, the proportion of mesenchymal-like cells was associated with increased metastatic potential. Similar associations were observed for the group of TNBC basal tumors which showed an overall higher fraction of mesenchymal-like cells.

## EMP is a continuum of cell states with intermediate EMP cells expressing distinct marker genes

Studies suggest that both mesenchymal and epithelial functions are necessary for the metastatic cascade (44 Therefore, the intermediate EMP cells were of special interest as they may represent cells with both epithelial and mesenchymal capabilities and a high degree of plasticity and therefore might contribute to the pool of cells that are more likely to metastasize (45, 46). However, the identified intermediate EMP cells, which expressed both epithelial and mesenchymal signatures at similar levels (Figure 4E), were present in every tumor although their abundance did not correlate with metastatic potential (Figure 4F). Intermediate EMP cells expressed core transcription factors (TFs) promoting EMT such as SNAI2, TWIST1, ZEB1 and ZEB2 (13) (upper panels of Figure 4G and Supplementary Figure S4I) at higher levels than epithelial-like cells but lower than mesenchymal-like cells highlighting their intermediate character. Moreover, the fraction of cells expressing these TFs also increased from epithelial-like to intermediate EMP to mesenchymal-like cells (lower panels of Figure 4G and Supplementary Figure S4I). To further characterize this intermediate EMP cell state we performed a more comprehensive differential gene expression analysis between the three EMP cell states in both datasets (Smart-Seq2 and MULTI-Seq) and identified genes upregulated in epithelial-like, intermediate EMP and mesenchymal-like cells (Figure 4H, I, Supplementary Table 7, Supplementary Table 8). For each EMP cell state, we focused on marker genes that were shared between the two datasets (Figure 4J). Surprisingly, only 13% (40/303, MULTI-Seq) – 18% (56/303, Smart-Seq2) of DEGs were shared with the published markers (43) that were used to classify the three EMP states. Most identified DEGs were exclusive to one or both of our datasets and were not found in the set of published markers (Supplementary Figure S4J). Genes shared across all three sets included common mesenchymal (e.g. VIM, BGN, SNAI2, LOX) and epithelial (e.g. KRT18, KRT8) markers; whereas other broadly used markers such as EPCAM, CDH1, CDH2 and FN1 were not included. Only 5 intermediate EMP cell marker genes were shared between our datasets (Figure 4J). The expression of all 5 intermediate EMP markers (CRYAB, KRT15, S100A2, CD24, CALML5) peaked in intermediate EMP cells and decreased in epithelial-like and mesenchymal-like cells (Figure 5A, Supplementary Figure S5A).

**Figure 5.**
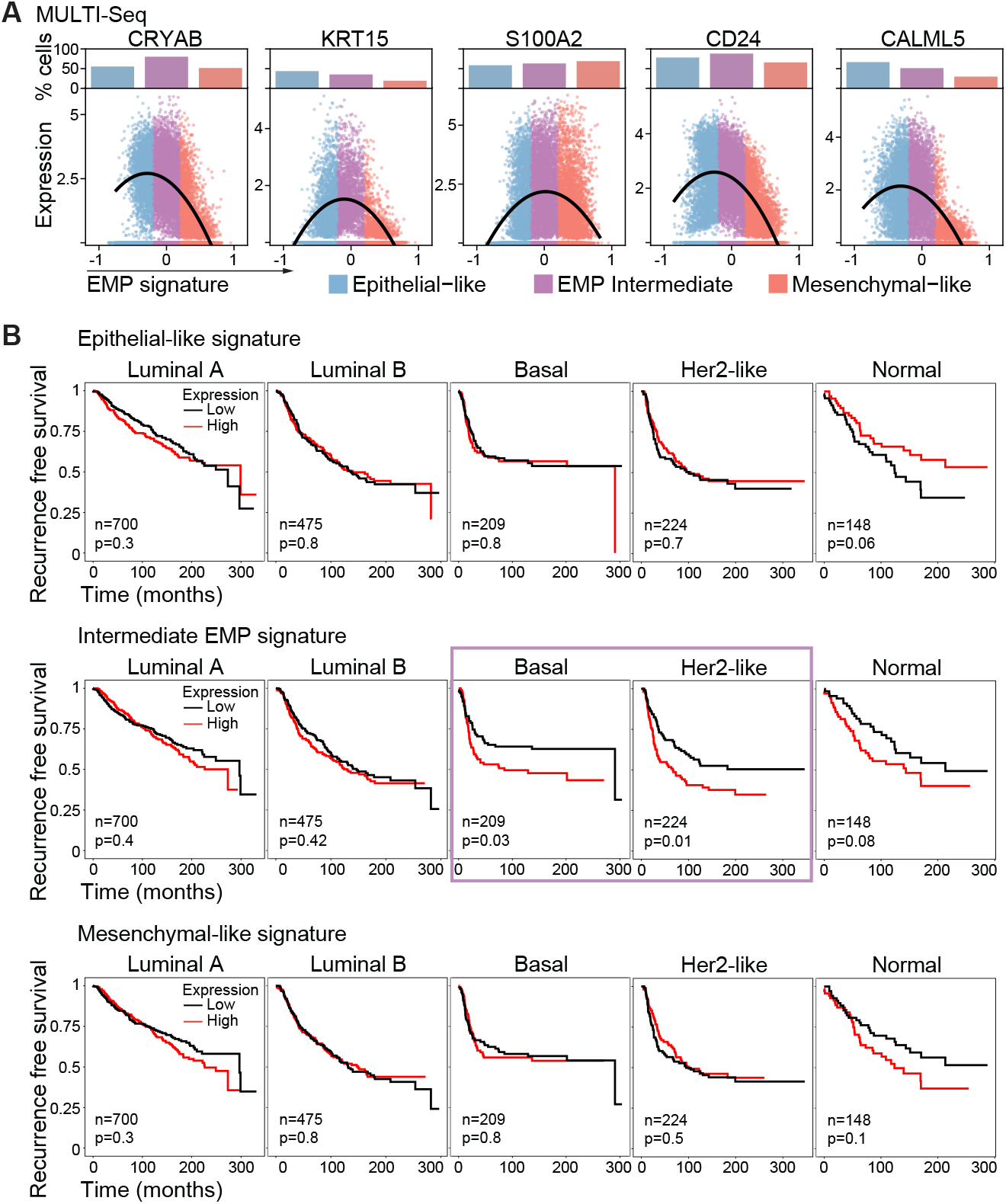
Intermediate EMP cell markers were correlated with patient outcome. (**A**) Scatter plots show the expression of indicated genes ordered by increasing EMP signature expression. Dots show the expression for individual cells and lines show smoothed expression of expressing cells. Bar charts on top show the proportion of positive expressing cells for the three EMP cell states (blue=epithelial-like, purple=intermediate EMP, red=mesenchymal-like cells). Showing the MULTI-Seq dataset. (**B**) Recurrence-free survival of BC patients (METABRIC) separated by PAM50 BC subtype using the mean expression of the epithelial-like (top panel), intermediate EMP (middle panel), and mesenchymal-like signatures (lower panel). The number of patients (n) and p-value (p) are shown. The purple box indicates a significant p-value.

The five markers of the intermediate EMP cell state have been previously implicated in EMT, cancer stemness, and metastasis pathways. For example, we identified the cell surface protein CD24, for which there are conflicting results as to its role in tumor progression. For example, whereas CD24^*−/*low^/CD44^+^ have been shown to initiate breast tumors in NOD/scid mice (47), other studies reported that high CD24 expression increased metastasis (48) and that CD24^+^/CD90^+^ cells initiate metastases that display a mesenchymal phenotype (49, 50). CD24^+^/CD44^+^ cells were shown to be plastic and express epithelial and mesenchymal markers forming mammospheres more efficiently than epithelial-like CD24^+^/CD44^*−*^ or mesenchymal-like CD24^*−*^/CD44^+^ cells (51–54). Interestingly, a mixture of CD24^+^/CD44^*−*^ and CD24^*−*^/CD44^+^ was more efficient in mammosphere formation than either population alone (52). Collectively, these data suggest that CD24 could indeed mark a plastic intermediate EMP cell state with potential stem-like properties and that cooperativity may exist between cell populations with different EMP characteristics.

Another identified intermediate EMP marker, CRYAB, encodes the small heat shock protein α-basic-crystallin (αB-crystallin), which protects cells from apoptosis by inhibiting caspase-3 activation under various stress conditions such as oxidative stress (55, 56). Importantly, αB-crystallin confers anoikis resistance and thereby enables metastatic dissemination (57). CRYAB is overexpressed in various tumors (58, 59) including the basal BC subtype (60) and has been associated with poor patient outcomes and metastasis (61–64). CRYAB expression levels in hepatocellular carcinoma cell lines were accompanied by EMT marker expression indicating that CRYAB could promote a mesenchymal phenotype (59). Importantly, a study investigating brain metastasis found that CRYAB is expressed in a small non-proliferative metastatic cell population and might be required for the survival of single metastatic cells and micrometastasis (65). Additionally, CRYAB is highly expressed in dormant micrometastasis in the lung compared to proliferative macrometastasis (65, 66). These combined features could also indicate an intermediate EMP cell state, although not explored in these studies.

Additional intermediate EMP markers were KRT15, CALML5 and S100A2. KRT15, an intermediate filament protein belonging to the epithelial Keratin Type I family, has been suggested to be an epidermal stem cell marker (67). One study reported that high KRT15 expression correlated with better outcomes for BC patients (68); whereas others reported that KRT15 was upregulated in advanced stage BC and BC with high-relapse risk (69, 70), as well as being associated with poor prognosis in other types of cancers (71, 72). CALML5 (Calmodulin-Like Protein 5) is a calcium-binding protein that is predominantly expressed in keratinocytes. It has recently been shown that calmodulin may mediate the induction of a partial EMP cell state (measured by loss of E-cadherin surface expression and a partial upregulation of mesenchymal markers) through calcium signaling (73). Also involved in calcium signaling is S100A2, which belongs to the S100 protein family that can both bind calcium and function extracellularly. S100A2 is deregulated in cancers suggesting both tumor-promoting and suppressing roles (74–76) and may also have dual roles with regard to EMT. S100A2 was shown to be regulated by TGF-β1 (77) and partially mediate TGF-β1-induced EMT (78, 79) but seems to repress EMT in other contexts (80).

Recent studies describing the existence of intermediate EMP cells have associated these with an increased ability to form metastases after tail vein injection using genetically engineered mouse models of skin squamous cell carcinoma (45). Here, we identified 5 novel markers, CD24, CRYAB, KR1T15, S100A2, and CALML5, that are expressed by human intermediate EMP cells in BC *in vivo* (Figure 5A, Supplementary Figure S5A). These genes could serve as biomarkers to identify BC patients with an increased proportion of potentially more aggressive tumor cells. To test the clinical significance of our findings we analyzed two BC gene expression datasets. In the first dataset (35), patients across different BC subtypes that showed a high expression of the epithelial-like gene signature had a better RFS, whereas patients with a high expression of the intermediate EMP or mesenchymal-like gene signature showed worse RFS (Supplementary Figure S5B). In an independent dataset (METABRIC (81)), intermediate EMP cell gene signature showed a BC subtype-dependent correlation with patient-related outcome (Figure 5B). Whereas luminal tumors did not show a correlation, high expression of the intermediate EMP cell signature in patients with basal and Her2-like classified BC showed worse RFS. These subtypes also showed the worst outcomes and resistance to therapy compared to the other subtypes (82).

Taken together, we identified cells co-expressing epithelial and mesenchymal markers that belong to an intermediate EMP cell state. These intermediate EMP cells were present in primary tumors and metastases of all tumor models studied and were characterized by low expression of EMT-associated TFs. Specific marker genes could identify intermediate EMP cells and a high expression of these markers was associated with worse patient-related outcomes. These novel intermediate EMP cell marker genes could serve as targets to block the dynamic process of EMP by directly targeting the potentially most plastic cells and thereby interfering with the metastatic cascade.

## Discussion

Metastasis is responsible for the majority of cancer-related deaths but the underlying processes that drive metastasis are not fully elucidated. Recent advances in single-cell biology shed light on the profound heterogeneity of tumors between and within patients that likely contributes to the complexity of the metastatic phenotype. By analyzing matched primary tumor and metastatic cells we found significant differences in the transcriptional profiles of metastatic cells compared to their primary tumor of origin in distinct BC subtypes that showed strong patient-to-patient variability. Specifically, we found that primary tumor cells consistently upregulated genes involved in hypoxia, glycolysis and other metabolic pathways across all tumor models. Moreover, metastatic cells frequently upregulated genes involved in cytoskeleton assembly, cell motility, cellular stress and immune response signaling. These transcriptional differences presumably are necessary to acquire traits for dissemination and are a result of the adaptations to different environments. A better understanding of this observed heterogeneity and the transcriptional differences between primary tumor and metastatic cells could have implications for therapy response.

One of these cellular traits is EMT, which has long been suggested as being an important driver of metastasis. Our current work highlights the complexity of the process (i.e. EMP) and its associated cell states. Recent research suggests that epithelial and mesenchymal cell states are the edges of a wider dynamic continuum of EMP including intermediate cell states (13, 83). However, these intermediate EMP cells remain poorly characterized. Here, we report that EMP is a dominant feature of tumor heterogeneity observed in different human BC tumors *in vivo*. We identified epithelial- and mesenchymal-like cells as well as intermediate EMP cells that surprisingly co-exist in every tumor. Intermediate EMP cells (described previously also as partial-EMT, hybrid-EMT, or EMT-transition cells) have recently been reported to exhibit the greatest metastatic potential when compared to mesenchymal or epithelial cells using tail vein injection of skin squamous carcinoma cells or orthotopic injection of highly metastatic pancreatic ductal adenocarcinoma cells, both derived from genetic mouse models (45, 46). On the contrary, we did not observe a correlation between the abundance of intermediate EMP cells in tumors and their metastatic potential. Indeed, we found that intermediate EMP cells were also present in very poorly metastatic tumor models. Instead, we found that a stronger mesenchymal phenotype (high EMP signature) and a higher proportion of mesenchymal-like cells correlated with the metastatic potential and this correlation could be further influenced by the BC subtype. Overall, our data suggest that the propensity to metastasize is a function of the entire tumor cell population, as opposed to the presence or absence of a ‘rogue’ and a potentially small subset of cells. It will be important for future studies to investigate the interactions of a heterogeneous tumor cell population with non-malignant cells of the environment and their involvement in the metastatic process.

Notably, our findings build upon, but do not necessarily contradict, the recently described importance of intermediate EMP cells for metastasis. Potentially, a higher proportion of epithelial-like cells may prevent intermediate EMP cells from metastasizing. Conversely, a higher proportion of mesenchymal cells may support the metastatic capabilities of intermediate EMP cells. One example of this proposed cooperativity between different EMP cells states is the observation that an admix culture of (epithelial-like) CD24^+^/CD44^*−*^ and (mesenchymal-like) CD24^*−*^/CD44^+^ immortalized normal human mammary epithelial (HMLER) cells was more efficient in mammosphere formation than either population alone (52). Thus, one hypothesis is that the critical factor determining the metastatic potential of a tumor is a combination of its composition of cells with varying EMP states and the level of cooperativity between them. The observation that metastasis can have polyclonal origins (84) and that circulating tumor cell (CTC) clusters are more effective in metastasis formation than individual CTCs (85) supports our idea that cooperativity between different EMP cell states may result in more effective metastasis formation.

EMP is likely a highly context and tumor type-specific process involving different signaling (6). Although the presence or proportion of intermediate EMP cells was not correlated with more metastasis in our models, we did find that high expression of intermediate EMP marker genes was associated with poorer outcomes in a subset of BC patients whereas a mesenchymal or epithelial gene expression did not show a subtype-specific correlation. This observation not only highlights the potential importance of the intermediate EMP cell state for patient outcomes but also indicates that the EMP process and its involvement in metastatic disease might be subtype-specific. Other markers of intermediate EMP cell states have been identified and linked to tumorigenesis, metastasis and stemness (such as CD104, EPCAM^*−*^/CD106^+^, ALDH1) (45, 54, 86, 87). These and our study highlight the need for a deeper understanding of the involvement of the intermediate EMP cell state in metastasis and its potential spatio-temporal context specify (23).

Surprisingly, there seems to be a predetermined, intrinsic equilibrium of EMP cell states within the tumor that appears to be independent of extrinsic signals and microenvironmental adaptations. Thus, primary tumor and metastatic cells exhibit very similar levels of the EMP signature expression despite showing remarkable differences in their overall transcriptomes. Sustaining plasticity and reversing the mesenchymal into a more epithelial-like cell state (MET) is proposed to occur during the formation of overt metastasis (88, 89). In this context, we would expect to detect more mesenchymal-like cells isolated from micrometastases and more epithelial-like phenotypes in macrometastases. Based on our histological characterization, poorly metastatic models show primarily micrometastases but also a few intermediate sized foci and potentially even rare macrometastases. Since metastatic cells were isolated from whole lung tissue, we were unable to distinguish whether cells obtained from poorly metastatic models were associated with intermediate-sized foci or (very rare) macrometastases or compare the transcriptome of micro- and macrometastases. Nonetheless, our single-cell analysis of primary tumor and metastasis revealed that EMP represents a continuum during spontaneous metastasis of a large panel of patient-derived breast tumors. Recent technology developments in spatial transcriptomics and multiplexed antibody-based imaging will be perfectly suited for future studies to investigate the dynamics of the various EMP cell states as metastatic tumors form.

## ACKNOWLEDGMENTS

We thank S. Schmid, H. Goodarzi, L. Murrow, Z. Gardell and H. Kortbai and all members of the Werb, Goga and Gartner labs for their valuable discussions and input. We thank M. Owyong, A. Abisoye-Ogunniyan, N. Ataii, and K. Salari for their technical assistance. We thank M.T. Lewis, L.E. Dobrolecki, and A. Welm for sharing their PDX models. We thank the UCSF flow core for their assistance, in particular A. Carlos, S. Kraus, and V. Nguyen. UCSF flow core is supported by RRID:SCR_018206 and DRC Center Grant NIH P30 DK063720. We thank E. Chow for sequencing assistance. We thank the UCSF Cancer Center Tissue Core. This study was supported by funds from EMBO long-term post-doctoral fellowship (EMBO ALTF 159-2017 to JW), Program for Breakthrough Biomedical Research Award (to JW), ImmunoX Bakar Trainee Momentum Award (to JW), Mark foundation (Endeavor grant to AG), Gazarian Foundation (to AG), Breast Cancer Research Foundation (to HR, AG), National Institutes of Health (U01 CA199315 to ZW, JW, AG), US National Institutes of Health 1R01CA223817 (AG), and the Chan Zuckerberg Biohub.

## AUTHOR CONTRIBUTIONS

JW conceptualized the study. WT, JW, CSM, MYH, DS, AA analyzed data. JW, CMD, SH, AN, VS, EA performed animal studies. JW, AA, JHH, WT, CMD, JVL performed tissue processing. JW, CSM prepared MULTI-Seq libraries. WT, MB, JW prepared Smart-Seq2 libraries. AD, NN performed sequencing. JW, CMD, AA, LA, SH performed tissue stainings. JW, SDu and SYL performed tissue imaging and analysis. JW, WT wrote the manuscript. SDa, MYH, AOP, AG, CSM edited the manuscript. SDa, AOP, AG, ZG, ZW provided guidance and funding.

## Materials and Methods

### Animal experiments

Fresh primary breast tumor samples were obtained from the Cooperative Human Tissue Network (CHTN) in accordance with the Institutional Review Boards’ approval. Tissues were received as de-identified samples and all subjects provided written informed consent. Medical reports were obtained without personally identifiable information. The UCSF Institutional Animal Care and Use Committee (IACUC) reviewed and approved all animal experiments. Tumor tissues were cut into 1 mm thick chunks and orthotopically transplanted into cleared mammary fat pats of 4-week-old NOD-SCID gamma mice to generate novel PDX models (J53353, J2036, and J55454, Supplementary Table 1). Established PDX lines were transplanted in the same way and as previously described (25, 28). Once palpable, tumors were measured 2×/week using a caliper to monitor growth kinetics. Tumor volume was calculated using following formula: 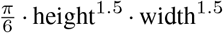. Unless otherwise noted, all PDX animals were euthanized at the endpoint, when the primary tumor reached 2.5 cm in diameter. In resection experiments, tumors were surgically removed at 1.0–2.0 cm in diameter. Resected animals were allowed to grow metastases until endpoint was reached (2.5 cm diameter of recurrent tumor). At endpoint, primary tumor and metastatic lungs were harvested, cut in small chunks and cryopreserved using Recovery Cell Culture Freezing Medium (Thermo Fisher, 12648010) and stored in liquid nitrogen until further analysis.

### Histology and tissue staining

For each PDX animal, after dissection, the middle and postcaval lobes of the right lung were fixed in 4% PFA overnight and processed for paraffin embedding. For histological analysis, tissue sections were stained with haematoxylin and eosin using standard protocols. Tissue slides were scanned (Zeiss Axio ScanZ.1) and images were analyzed using QuPath. Metastatic foci were easily identified by a larger nuclei/cytoplasm ratio. Micrometastases were defined as < 10 tumor cells, intermediate metastatic foci 10–100 cells and macrometastases >100 cells. Number and area of metastatic foci and total tissue area were determined.

### Lysis plate preparation

Lysis plates were prepared by dispensing 0.4 µL lysis buffer (0.5 U Recombinant RNase Inhibitor (Takara Bio, 2313B), 0.0625% Triton™ X-100 (Sigma, 93443-100ML), 3.125 mM dNTP mix (Thermo Fisher, R0193), 3.125 µM Oligo-dT30VN (IDT, 5^*′*^-AAGCAGTGGTATCAACGCAGAGTACT30VN-3^*′*^) and 1:600,000 ERCC RNA spike-in mix (Thermo Fisher, 4456740)) into 384-well hard-shell PCR plates (Biorad HSP3901) using a Dragonfly liquid handler (STP Labtech). All plates were then spun down for 1 min at 3220×g and snap-frozen on dry ice. Plates were stored at -80 °C until used for sorting.

### Sample preparation and FACS sorting

Primary tumor and metastatic lung tissues were processed, stained, MULTI-Seq labeled and FACS sorted as previously described (36). In brief, tissues were thawed, dissociated in digestion media containing 50 µg/ml Liberase TL (Sigma-Aldrich) and 2·10^4^ U/ml DNase I (Sigma-Aldrich) in DMEM/F12 (Gibco) using standard GentleMacs (37C_m_LDK_1, 37_m_TDK1) protocols. Washed and filtered single-cell suspension were stained with viability dye (Zombie NIR, 1:500, BioLegend, no. 423105), blocked with Fc-block (1:200, Tonbo, 70-0161-U500), and with LIN (anti-mouse TER119-FITC, Thermo Fisher, 11-5921-82; anti-mouse CD31-FITC, Thermo Fisher, 11-0311-85; anti-mouse CD45-BV450, Tonbo, 75-0451-U100; anti-mouse MHC-I-APC, eBioscience, 17-5999-82) and anti-human CD298 (PE, BioLegend, 341704). For the Smart-Seq2 experiments, live, LIN^*−*^/hCD298^+^ primary tumor and metastatic cells were sorted directly into cooled lysis plates and snap-frozen until library preparation. If multiple plates were sorted from one PDX model, each plate contained half primary tumor and metastatic cells to avoid plate-specific batch effects. For the MULTI-Seq experiments, MULTI-Seq LMO barcode anchor and co-anchor were used at a final concentration of 2.5 µM directly after antibody staining before FACS sorting as described previously (36). For one experiment (PDX1) we used sets of three unique MULTI-seq barcodes/sample. After sorting, enriched live, LIN^*−*^/hCD298^+^ cells were pooled and loaded into 10x microfluidics lanes at an average loading concentration of about 30,000 cells/lane.

### cDNA synthesis and library preparation

cDNA synthesis was performed using the Smart-seq2 protocol (90–92). Briefly, 384-well plates containing single-cell lysates were thawed on ice followed by first-strand synthesis. 0.6 µL of reaction mix (16.7 U/µl SMARTScribe Reverse Transcriptase (Takara Bio, 639538), 1.67 U/µl Recombinant RNase Inhibitor (Takara Bio, 2313B), 1.67X First-Strand Buffer (Takara Bio, 639538), 1.67 µM TSO (Exiqon, 5^*′*^-AAGCAGTGGTATCAACGCAGACTACATrGrG+G-3^*′*^), 8.33 mM DTT (Bioworld, 40420001-1), 1.67 M Betaine (Sigma, B0300-5VL), and 10 mM MgCl2 (Sigma, M1028-10×1ML)) was added to each well using a Dragonfly liquid handler (STP Labtech). Reverse transcription was carried out by incubating wells on a ProFlex 2×384 thermal-cycler (Thermo Fisher) at 42 °C for 90 min and stopped by heating at 70 °C for 5 min. Subsequently, 1.5 µL of PCR mix (1.67X KAPA HiFi HotStart ReadyMix (Kapa Biosystems, KK2602), 0.17 µM IS PCR primer (IDT, 5^*′*^-AAGCAGTGGTATCAACGCAGAGT-3^*′*^), and 0.038 U/µl Lambda Exonuclease (NEB, M0262L)) was added to each well with a Dragonfly liquid handler (STP Labtech), and second strand synthesis was performed on a ProFlex 2×384 thermal-cycler by using the following program: 1. 37 °C for 30 min, 2. 95°C for 3 min, 3. 23 cycles of 98 °C for 20 s, 67 °C for 15 s, and 72 °C for 4 min, and 4. 72 °C for 5 min. The amplified product was diluted 1:10 with 10 mM Tris-HCl (Thermo Fisher, 15568025). 0.6 µL of diluted product was transferred to a new 384-well plate using the Viaflow 384 channel pipette (Integra). Illumina sequencing libraries were prepared using a library preparation protocol modified from previously reported tagmentation-based protocols (93, 94). Briefly, tagmentation was carried out by mixing each well with 1 uL of 1.6x Homebrew Tn5 Tagmentation Buffer and 0.2 uL of homebrew Tn5 enzyme, then incubated at 55 °C for 3 min. The reaction was stopped by adding 0.4 µl 0.1% sodium dodecyl sulfate (Fisher Scientific, BP166-500) and centrifuging at room temperature at 3,220g for 5 min. Indexing PCR reactions were performed by adding 0.4 µL of 5 µM i5 indexing primer, 0.4 µL of 5 µM i7 indexing primer, and 1.2 µL of Nextera NPM mix (Illumina). All reagents were dispensed with the Mosquito liquid handlers (STP Labtech). PCR amplification was carried out on a ProFlex 2×384 thermal cycler using the following program: 1. 72 °C for 3 min, 2. 95°C for 30 s, 3. 12 cycles of 98 °C for 10 s, 67 °C for 30 s, and 72 °C for 1 min, and 4. 72 °C for 5 min.

### Library sequencing

Following library preparation, wells of each library plate were pooled using a Mosquito liquid handler (STP Labtech). Pooling was followed by two purifications using 0.7x AMPure beads (Fisher, A63881). Library quality was assessed using high sensitivity capillary electrophoresis on a Tapestation (Agilent), and libraries were quantified by qPCR (Kapa Biosystems, KK4923) on a CFX96 Touch Real-Time PCR Detection System (Biorad). Plate pools were normalized to 2 nM and equal volumes from library plates were mixed together to make the sequencing sample pool. Sequencing libraries from 384-well plates Libraries were sequenced on the NextSeq or NovaSeq 6000 Sequencing System (Illumina) using 2×100 bp paired-end reads and 2×12 bp index reads. NextSeq runs used high output kits, whereas NovaSeq runs used 300-cycle kit (Illumina, 20012860). PhiX control library was spiked in at ∼1%.

### Sequencing libraries from MULTI-seq

For the MULTI-Seq dataset, gene expression library preparation was performed using the v2 10x library kit with modifications as described previously to generate MULTI-seq libraries (36).

### Data extraction

For Smart-Seq2, sequences from the NovaSeq or NextSeq were de-multiplexed using bcl2fastq v.2.19.0.316. Reads were aligned to the gencode V30 genome using STAR v.2.5.2b with parameters TK. Gene counts were produced using HTSEQ v.0.6.1p1 with default parameters, except ‘stranded’ was set to ‘false’, and ‘mode’ was set to ‘intersection-nonempty’. For MULTI-Seq, sequences from the microfluidic droplet platform were de-multiplexed and aligned using CellRanger v.5.0.1, available from 10x Genomics with default parameters.

### MULTI-Seq demultiplexing

MULTI-seq barcode FASTQs were converted to barcode UMI count matrices using the ‘MULTIseq.preProcess’ and ‘MULTIseq.align’ functions in the deMULTIplex R package (36) with default parameters. Notably, ‘PDX3’ FASTQs were randomly down-sampled to 10^8^ total reads prior to UMI count matrix conversion in order to minimize computation time. Next, since cells labeled with the same MULTI-seq barcodes were split across multiple 10x Genomics microfluidics lanes in each experiment, MULTI-seq UMI count matrices from each lane were concatenated (PDX1 and PDX3 matrices were concatenated separately) to maximize classification performance. Using these concatenated matrices, samples were then classified into sample groups using the deMULTIplex workflow desired previously (with semi-supervised negative-cell reclassification) (36). Notably, since samples in the PDX1 experiment were encoded by sets of three unique MULTI-seq barcodes, classification was performed on each cell’s median barcode count for each set. Moreover, cells with the top and bottom 5% of MULTI-seq barcode counts were masked during the initial classification workflow in the PDX1 data, and were reintroduced as ‘negatives’ during semi-supervised negative cell reclassification. Analogous barcode count-merging and outlier-masking were not necessary for the PDX3 data, which was classified successfully using the default deMULTIplex workflow.

### Data pre-processing

For Smart-seq 2 data, gene count tables were combined with the metadata variables using the Scanpy Python package version 1.8.1 (95). We removed the genes that were not expressed in at least 5 cells. Cells with less than 5,000 counts and 500 detected genes were removed. Additionally, we removed cells with more than 50% mitochondrial genes and 20% ERCC reads. The data were then normalized using size factor normalization such that every cell has 10,000 counts and log-transformed. We selected the top 2,000 genes with the highest standardized variance as the highly variable genes by using VST method from Seurat V3 (96), which is also implemented in Scanpy. Cell-cycle regression was performed after calculating the score of S and G2M phases for each cell (97). The data was then scaled to a maximum value of 10. We then computed principal component (PC) analysis, neighborhood graph and clustered the data using Louvain and Leiden methods. The data was visualized using UMAP projection.

For MULTI-seq data, for each 10x lane, we first removed the cells with less than 2,500 UMI and 250 genes and more than 50% mitochondria reads by using the Scanpy Python package version 1.8.1. In addition, in order to filter out reads from ambient RNA, we ran DecontX (98) separately for each 10x lane by using default parameters. Next, we re-filtered the dataset from every 10x run when cells did not contain a minimum number of genes (250), minimum of counts/UMIs (2,500), and/or having more than 50% mitochondria reads. The data were then further processed as described above for the Smart-Seq2 dataset. Cells were sample assigned using the MULTI-Seq demultiplexing result, thereby removing doublets but including unassigned ‘negative’ cells. In order to recover MULTI-Seq-unassigned ‘negative’ cells, we used DBSCAN clustering. Based on the results of the Smart-Seq2 data, cells from different PDX tumors would cluster distinct from each other in transcriptional space. Negative cells in one DBSCAN cluster were assigned as the same tumor sample as the majority of MULTI-Seq classified cells in that cluster. After completed cell assignment, all 10x runs were combined to one MULTI-Seq dataset and removed genes that were not expressed in at least 20 cells. We then performed normalization, log-transformation, finding highly variable genes, cell cycle regression, principal component analysis, UMAP dimension reduction, and Louvain and Leiden clustering as described for the Smart-Seq data.

### EMP scoring and classification

We used GSVA scoring with its default parameters to assign each cell an epithelial (E-score) and mesenchymal score (M-score) using epithelial and mesenchymal marker genes (43). The EMP-score for each cell is calculated by the sum of E-score and M-score for that cell. Cells with an EMP-score > 0.2 were classified as mesenchymal-like cells, cells with an EMP-score < -0.2 were classified as epithelial-like cells, and cells with an EMP-score between -0.2 and 0.2 were classified as EMP intermediate cells.

### Cell Phase Proportion Statistical Test

Cells were assigned in different cell cycle phases based on the cell cycle score calculated previously. Then cell phase proportions in each tumor were calculated in different groups in each category, such as EMP cell stage, sort, and metastatic potential group. Finally, Wilcoxon rank test was performed for comparing group to group in each category in each cell phase. The statistical tests were generated by using Seaborn (99) and Statannot (100) packages in Python.

### ROC Curve and AUC Value

The cells were first ordered by their PC2 value either in the whole SS2 dataset or in individual tumor models. The true and false-positive rates were calculated based on the cell’s label (primary tumor cell or metastatic cell) and PC2 value by using the “roc_curve” function from Scikit-learn (101). In addition, “roc_auc_score” from Scikit-learn was used to calculate the AUC value by using the true and false-positive rates.

### Differentially expressed genes

#### Identifying DEGs in primary tumor and metastatic cells

We performed differential expression analysis between primary tumor and metastatic cells in the entire dataset using the Seurat function FindMarkers using MAST (33) and the tumor model as the latent variable. In addition, we identified DEGs between primary tumor and metastatic cells for each tumor sample separately using the same Seurat function without setting the latent variable. Genes with p-values < 0.05 and log_2_ fold change > 0.5 were kept for further analysis. After filtering, we combined the DEGs from tumors within the same metastatic potential group. We included DEGs that are shared between at least two tumors in the same metastatic potential group.

#### Identifying gene signatures associated with metastatic potential

To identify genes in the primary tumor that are associated with metastatic potential we first removed metastatic cells from the data. Then, we used the Seurat function FindMarkeres using the MAST test and tumor model as the latent variable for identifying DEGs between one individual tumor and all tumors in the other metastatic groups. Genes were filtered based on p-values < 0.05 and log_2_ fold change > 0.5. After filtering, we combined the up-regulated gene lists from tumors within the same metastatic potential group. Signature genes related to metastatic potential were determined as genes that are shared between at least two tumors in the same metastatic potential group.

#### Identifying EMP marker genes

We identified EMP marker genes for each EMP category (epithelial-like, EMP intermediate, mesenchymal-like) using the Seurat function FindMarkers using the MAST test and tumor model as the latent variable. Genes were filtered based on p-values < 0.05 and log_2_ fold change > 0.5.

### Gene set enrichment analysis

To identify pathways that were enriched in primary tumor or metastatic cells, we used the fgsea package (102) with Hallmark and GO gene sets from MSigDB (103, 104). We examined pathways that were significantly enriched in at least four tumor models. Enriched pathways in highly and poorly metastatic signatures were identified with the online tool of MSigDB.

### Survival Analysis

For survival analysis, we used the KM-plotter (35) website’s breast cancer gene chip mRNA dataset. The mean expression of the signature genes were calculated for each sample in the dataset. The patient samples were separated based on the median of the mean expression in low and high expressing samples. Visualization was done using Lifelines Python package (105). The metastatic potential gene lists resulted from the overlap genes between the MULTI-seq and Smart-Seq2 datasets from each metastatic potential group. For EMP signature gene lists, the epithelial signature gene list and mesenchymal signature gene list were the overlapping genes from the top 100 differentially expressed genes in the MULTI-Seq and Smart-Seq2 datasets, and the intermediate EMP marker gene signature included the overlapping genes found in both MULTI-seq and Smart-Seq2 datasets. In addition, METABRIC (81) dataset was obtained from cBioPortal. We used “all_sample_Zscores” to create Kaplan-Meier survival plots (105) and perform logrank tests for each breast cancer subtype using the mean expression of the intermediate EMP marker genes.

## Supplemental Tables

- Table S1. PDX info
- Table S2. SS2 met vs. primary DEGs filtered (Tabs: global, HCI005, H3404, H4272, HCI009, HCI011, HCI001, H5097, J2036, J53353, HCI010)
- Table S3. Overlaped DEGs primary tumor vs. metastasis per metastatic potential group
- Table S4. MULTI primary tumor 1 vs. rest DEGs filtered (Tabs: HCI002_MULTI, J55454_MULTI, HCI005_MULTI, H4272_MULTI, HCI011_MULTI, HCI001_MULTI, H5097_MULTI, J2036_MULTI, J53353_MULTI, HCI010_MULTI)
- Table S5. SS2 primary tumor 1 vs. rest DEGs filtered (Tabs: J55454_SS2, H5471_SS2, HCI005_SS2, H3404_SS2, H4272_SS2, HCI009_SS2, HCI011_SS2, HCI001_SS2, H5097_SS2, J2036_SS2, J53353_SS2, HCI010_SS2)
- Table S6. Low, moderate, high met. signature
- Table S7. SS2 EMP DEGs filtered (Tabs: Epithelial-like, Mesenchymal, EMP intermediate)
- Table S8. MULTI EMP DEGs filtered (Tabs: Epithelial-like, Mesenchymal, EMP intermediate)

## Data and materials availability

Raw sequencing files are available at the NCBI BioProject number PRJNA847563. Raw and processed data have been deposited in NCBI’s Gene Expression Omnibus and are accessible through GEO Series accession number GSE210283. Processed data are available as h5ad files on figshare (https://figshare.com/s/328942c0b8dc9aa69be1 and https://figshare.com/s/b53f327a8b612a7b2eeb). Code is available on github https://github.com/czbiohub/scBC.

---

**Figure S1.**
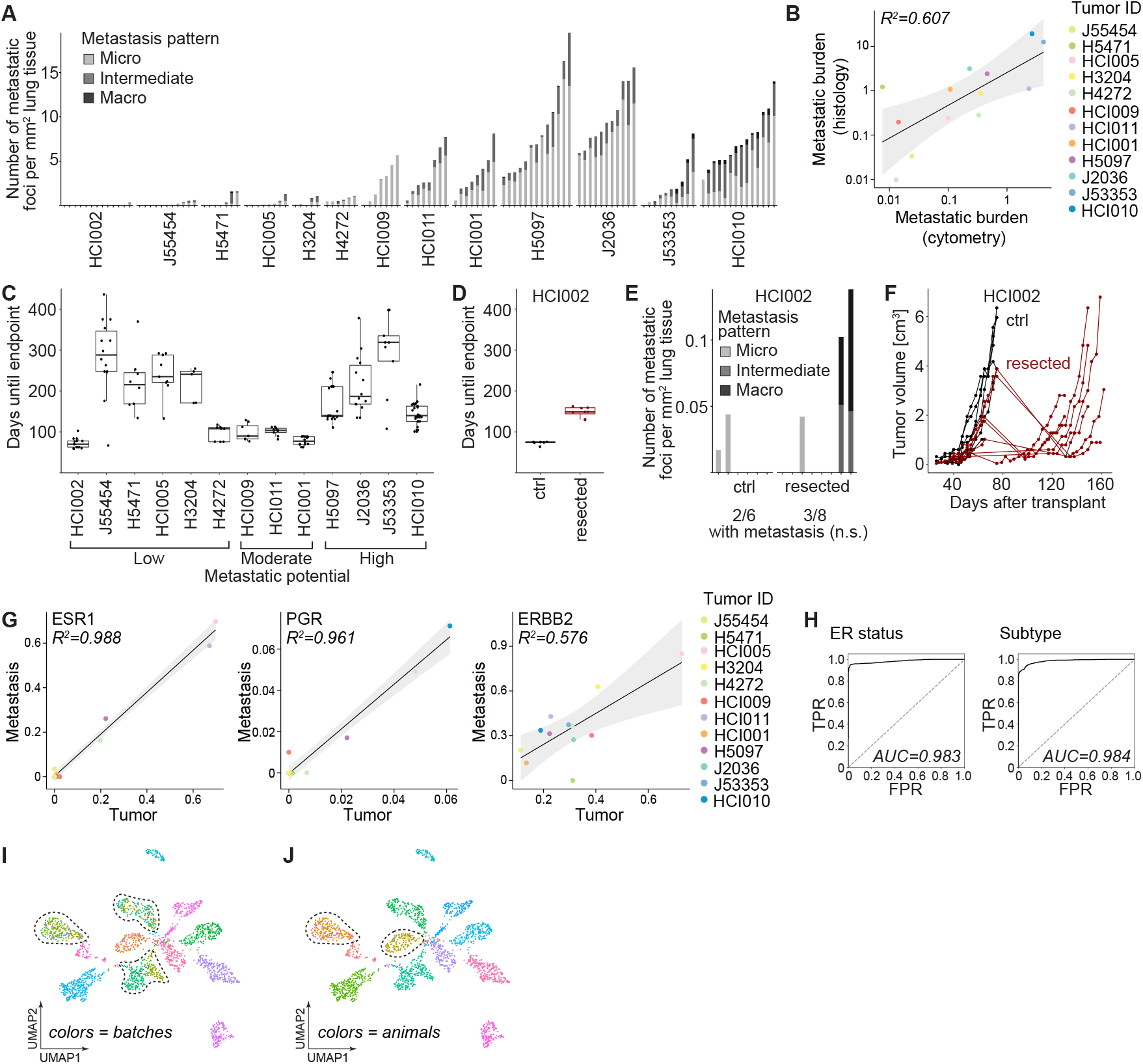
Biological characteristics of the PDX models. (**A**) Bar chart shows the number of metastatic foci per mm^2^ lung tissue area for individual animals ordered by the metastatic potential of the tumor models determined by histology. Each tick mark represents one animal. The size of metastatic foci is colored in shades of gray (micrometastasis < 10 cells, intermediate 10–100 cells and macrometastasis > 100 cells). (**B**) Scatter plot shows the correlation of mean metastatic burden assessed by histology (proportion of metastatic tissue area to total lung tissue area) and flow cytometry (proportion of metastatic cells to total live cells) colored by individual tumor models. Linear regression with 95% confidence intervals and Pearson correlation coefficient are shown. (**C**) Boxplot shows median days until endpoint (2.5 cm diameter of primary tumor) after tumor transplantation per tumor model ordered by the metastatic potential as determined in Figure 1B. (**D**) Boxplot shows median days until endpoint (2.5 cm diameter of the primary tumor or recurrent tumor) after tumor transplantation comparing HCI002 (black) and after HCI002 resection (red). (**E**) Bar chart shows the number of metastatic foci per mm^2^ lung tissue area for individual animals of HCI002 and resected HCI002 at endpoint. The size of metastatic foci is colored in shades of gray (micrometastasis < 10 cells, intermediate 10–100 cells and macrometastasis > 100 cells). Each tick mark represents one animal showing 2/6 (control) and 3/8 animals (resected) that developed metastases (p-value=0.872, Chi-Square test). (**F**) Spider plot shows tumor growth (volume in cm^3^) for each animal transplanted with HCI002 (black) or resected with subsequent recurrent tumor (red). (**G**) Scatterplots show the correlation of the mean expression of the indicated receptors in primary tumor and metastatic cells colored by individual tumor models. Linear regressions with 95% confidence intervals and Pearson correlation coefficients are shown. (**H**) ROC curves with the corresponding area under the curve (AUC) show PC1 categorized based on ER status (left) and BC subtype (right). (**I**) UMAP projection of single-cell transcriptomes color-coded by batch (individual plates). Dashed lines highlight clusters of cells from the same tumor model measured in multiple batches (technical replicates). (**J**) UMAP projection of single-cell transcriptomes color-coded by individual animals. Dashed lines highlight clusters of cells from the same tumor model retrieved from multiple animals (biological replicates).

**Figure S2.**
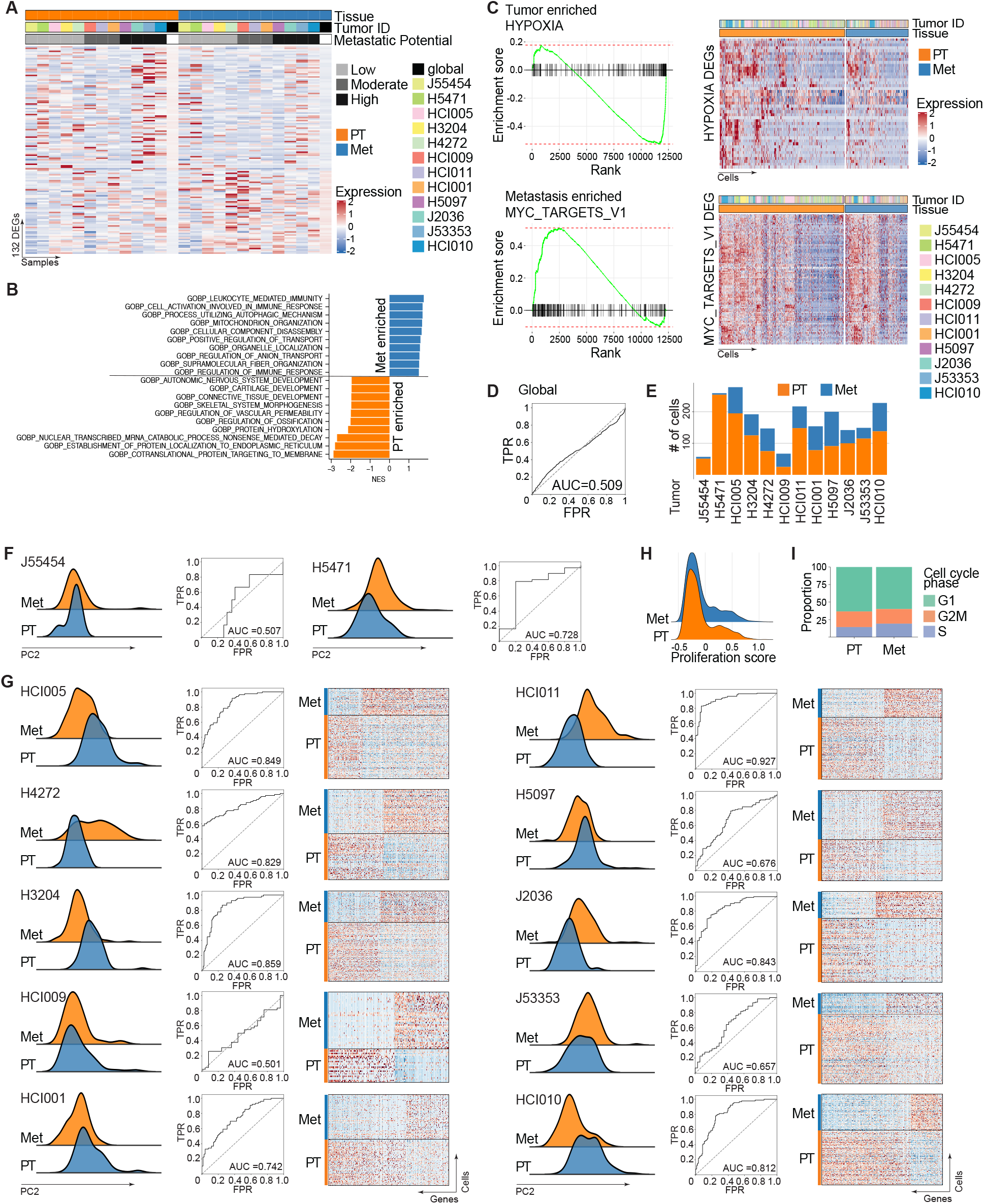
Differential gene expression between primary tumor and matched metastatic cells. (**A**) Heatmap shows mean expression per tumor model of DEGs between primary tumors and metastases. Annotations indicate tissue, tumor model, and metastatic potential. (**B**) Bar chart shows pathways enriched in primary tumors (negative NES, orange) and metastases (positive NES, blue) using GO biological pathways from MSigDB. (**C**) Enrichment plots show hypoxia as the top enriched pathway in primary tumors (top) and MYC targets as the top enriched pathway in metastases (bottom). Heatmaps show expression in single cells of DEGs associated with either hypoxia (top) or MYC targets (bottom). Annotations show tissue and tumor model. (**D**) ROC curve using PC2 coordinates to classify cells into either primary tumor or metastatic cells of all tumors grouped together (global) with depicted AUC. (**E**) Bar chart shows the number of primary tumor (orange) and metastatic cells (blue) for each tumor model. (**F**) Ridge plots show normalized cell counts along PC2 color-coded by primary tumor and metastasis for tumor models J55454 and H5471 without a sufficient number of metastatic cells and corresponding ROC curves (same as in (D)). (**G**) Ridge plots show normalized cell counts along PC2 color-coded by primary tumor and metastasis for individual tumor models with a sufficient number of metastatic cells and corresponding ROC curves of PC2 (same as in (D)). Clear separations in PC2 are reflected by AUC > 0.7 by ROC curve analysis. Heatmaps show the expression of DEGs between primary tumor and metastatic cells. (**H**) Ridge plot shows proliferation score for primary tumors and metastases. (**I**) Bar chart shows the proportion of cells in G1/G2M/S cell cycle phase for primary tumors and metastases. Not significant using Wilcoxon rank test.

**Figure S3.**
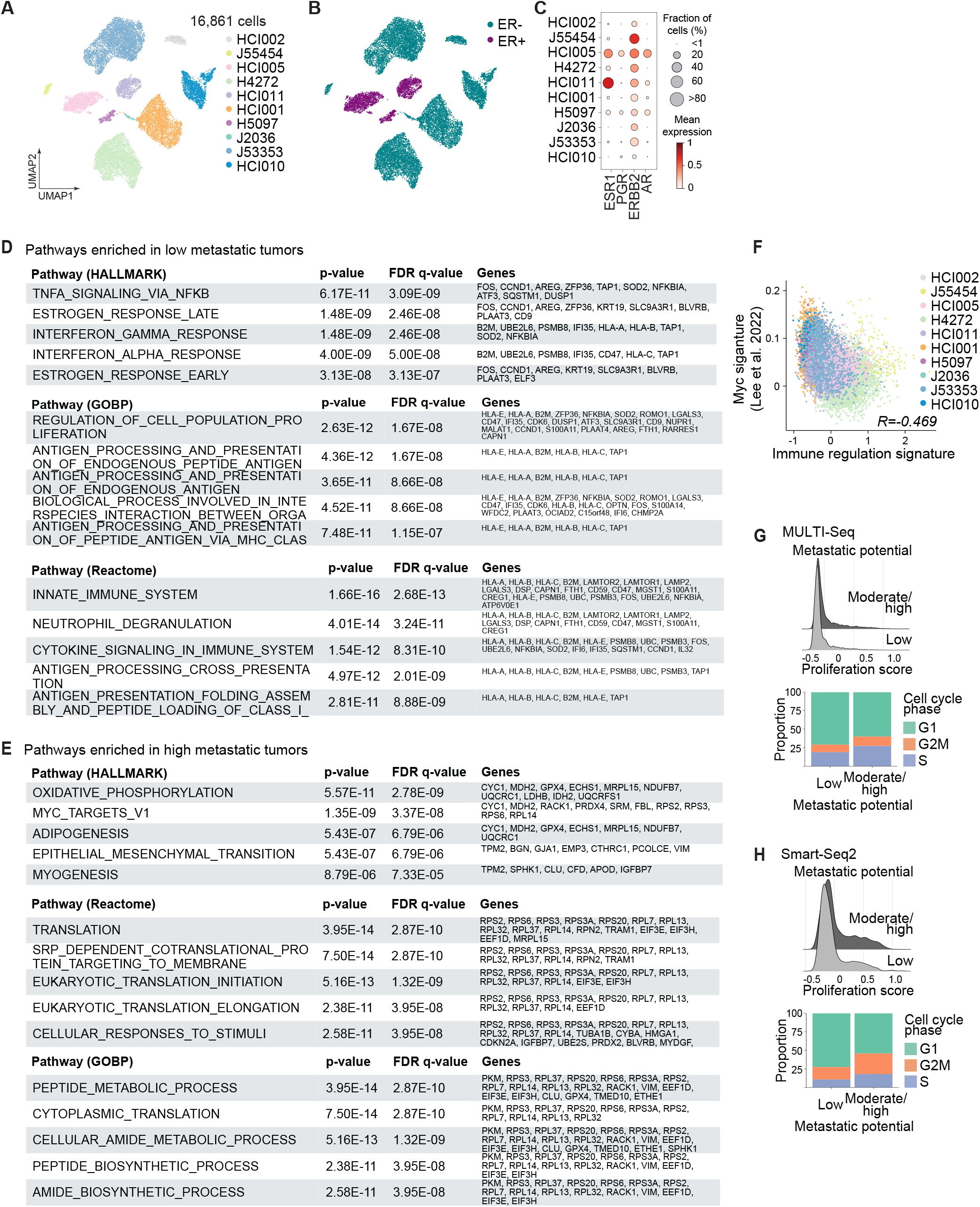
Characteristics of the metastatic signatures. (**A**) UMAP projection of single-cell transcriptomes color-coded by individual tumor models. (**B**) UMAP projection of single-cell transcriptomes color-coded by ER status. (**C**) Bubble plot shows the expression of receptors per tumor model. The size of dots indicates the fraction of cells expressing and the red color indicates gene expression. (**D**) Pathway enrichment of DEGs shared between poorly metastatic tumors. (**E**) Pathway enrichment of DEGs shared between highly metastatic tumors. (**F**) Scatterplot shows the correlation of MYC (42) and immune regulation signature expression colored by tumor model. Pearson correlation coefficient is shown. (**G**) Ridge plot shows proliferation score for primary tumor of low and moderate/high metastatic potential. The bar chart shows proportion of cells in different cell cycle phases for primary tumors of low and moderate/high metastatic potential. Showing MULTI-Seq dataset. Proportion changes are not significant using Wilcoxon rank test. (**H**) Same as in (G) for the Smart-Seq2 dataset. Not significant using Wilcoxon rank test.

**Figure S4.**
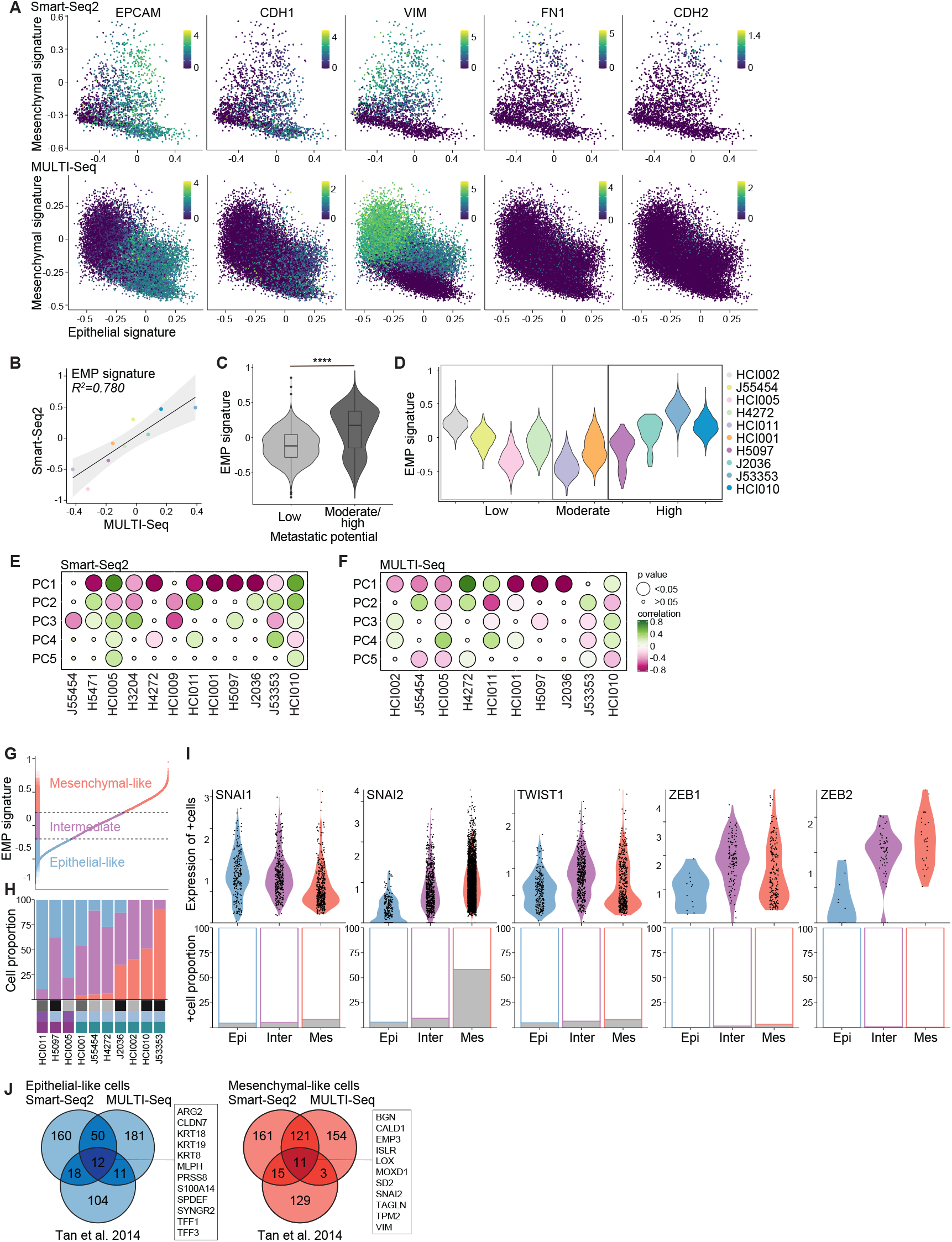
EMP is a key feature of tumor heterogeneity. (**A**) Scatter plots show mesenchymal against epithelial signatures for individual cells colored by the expression of indicated epithelial (EPCAM, CDH1) and mesenchymal markers (VIM, FN1, CDH2). The upper panels show Smart-Seq2 and the lower panels show MULTI-Seq datasets. The color scale indicates the magnitude of gene expression. (**B**) Scatter plot shows the correlation of the mean EMP signature expression per tumor model between Smart-Seq2 and MULTI-Seq datasets. Linear regression with 95% confidence intervals and Pearson correlation coefficient are shown. (**C**) Violin plot shows EMP signature expression of tumor models with low and intermediate/high metastatic potential using the MULTI-Seq dataset. Boxplot showing median, significance p<0.001 by Wilcox test. (**D**) Violin plot shows EMP signature expression per tumor model ordered by metastatic potential using the MULTI-Seq dataset. (**E**) Bubble plot shows the correlation of EMP signature with PCs 1-5 using Smart-Seq2 dataset. The color indicates positive (green) or negative (purple) correlation coefficient, larger circle indicates significant p-value < 0.05, small circle indicates no significant p-value > 0.05. (**F**) same as in (E) for the MULTI-Seq dataset. (**G**) Cells ranked by EMP signature defining three cell states: epithelial-like (blue), intermediate EMP (purple) and mesenchymal-like cells (red) using the MULTI-Seq dataset. (**H**) Bar chart shows the proportion of the three different EMP cell states per tumor model ranked by the increasing proportion of mesenchymal-like cells. Gray-scale boxes indicate the metastatic potential. Other annotations indicate ER status and BC subtype as in Figure 4F. Showing MULTI-Seq dataset. (**I**) Violin plots (top) show expression of EMT-associated TFs in expressing cells grouped by EMP cell states (Epi = epithelial-like, Inter = Intermediate EMP, Mes = mesenchymal-like cells). Bar charts (bottom) show the fraction of expressing cells in gray. Showing MULTI-Seq dataset. (**J**) Venn diagrams show overlaps of epithelial (blue, left panel) and mesenchymal markers (red, right panel) for Smart-Seq2, MULTI-Seq and Tan et al. 2014. Highlighted are genes shared between all three sets.

**Figure S5.**
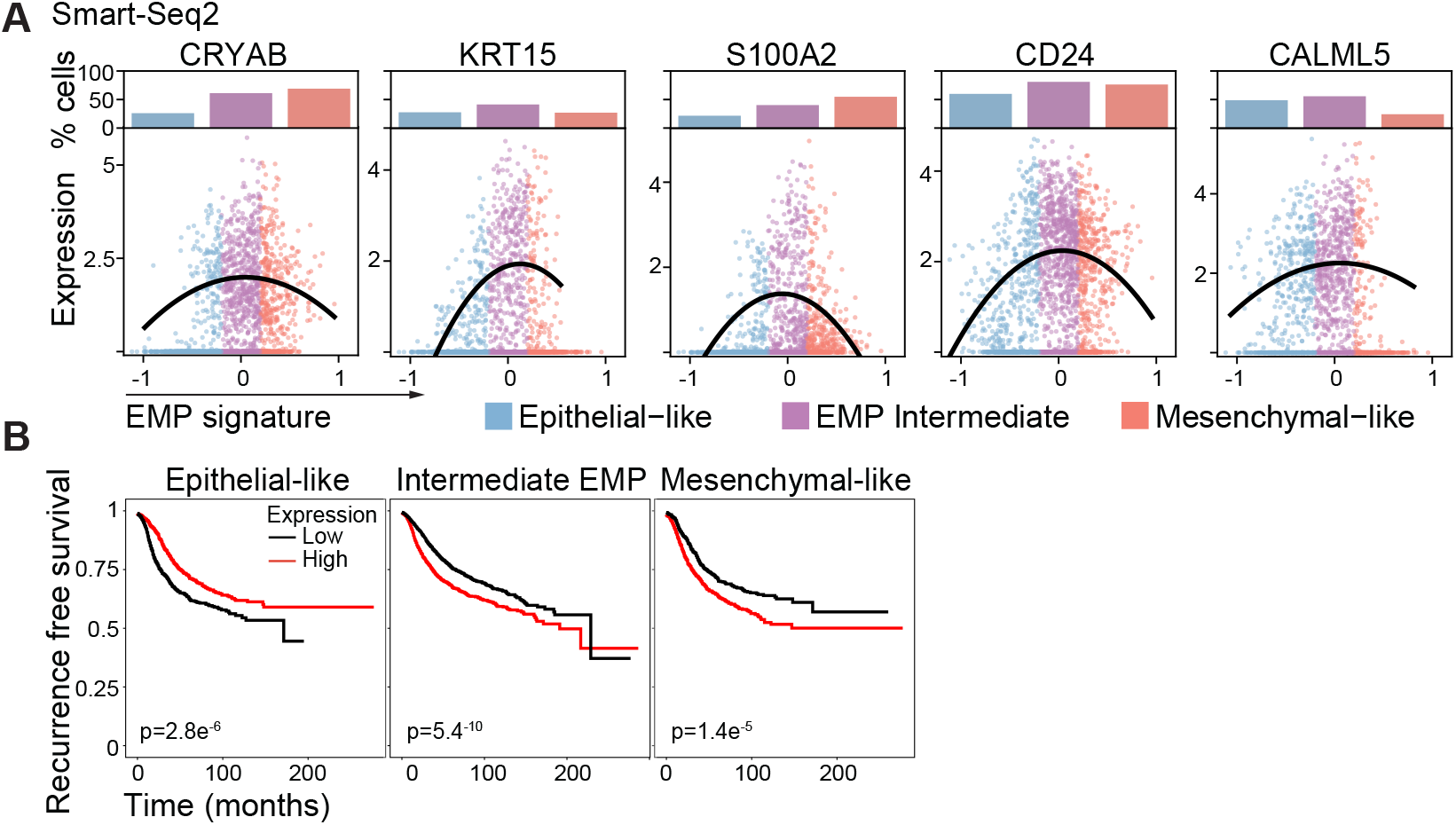
Intermediate EMP cell markers were correlated with patient outcome. (**A**) Scatter plots show the expression of indicated genes ordered by increasing EMP signature expression. Dots show expression for individual cells, lines show smoothed expression of expressing cells. Bar charts on top show the proportion of positive expressing cells for the three EMP cell states (blue=epithelial-like, purple=intermediate EMP, red=mesenchymal-like cells). Showing the Smart-Seq2 dataset. (**B**) Recurrence-free survival of BC patients using the mean expression of the overlapped genes for each EMP cell state (generated with KM-plotter (35)).

## Notes

### Competing Interest Statement

The authors have declared no competing interest.

